# Molecular dynamics guided engineering of *Aequorea victoria* Green Fluorescent Protein chromophore interactions generates a brighter variant with improved photobleaching resistance

**DOI:** 10.1101/2025.01.08.631926

**Authors:** Rochelle D. Ahmed, W. David Jamieson, Danoo Vitsupakon, Athena Zitti, Kai A. Pawson, Oliver K. Castell, Peter D. Watson, D. Dafydd Jones

**Author notes:** These authors contributed equally and are ordered alphabetically based on surname (Rochelle D. Ahmed, W. David Jamieson, Danoo Vitsupakon, Athena Zitti). Corresponding author: D. Dafydd Jones, School of Biosciences, Sir Martin Evans Building, Cardiff University, Cardiff, CF10 3AX, UK.

## Abstract

Fluorescent proteins (FPs) are a crucial tool for cell imaging, but with developments in fluorescence microscopy and researcher requirements there is still a need to develop brighter versions that remain fluorescent for longer. Using short time-scale molecular dynamics-based modelling to predict changes in local chromophore interaction networks and solvation, we constructed an *Aequorea victoria* GFP (avGFP) variant called YuzuFP that is 1.5 times brighter than the starting superfolding variant (sfGFP) with a near 3-fold increased resistance to photobleaching *in situ*. YuzuFP contained a single mutation that replaces the chromophore interacting residue H148 with a serine. Longer time scale molecular dynamics revealed the likely mechanism of action is S148 makes more persistent polar interactions with the chromophore phenol group and increases the residency time of an important water molecule. As demonstrated by live cell imaging, YuzuFP not only offers a timely upgrade as a useful green-yellow avGFP for cell imaging applications over longer timescales, but it also provides a basic scaffold for future avGFP engineering efforts.

## Introduction

Fluorescent proteins (FPs) are an indispensable tool in modern cell biology ^1–4^, with protein engineering playing a pivotal role in generating useful variants from a relatively small pool of non-optimal natural starting points. However, there is still a continuing need to generate new FPs to match the improvements and requirements in cell imaging. Such parameters include improved brightness, resistance to photobleaching, longer “on” times and longer fluorescence lifetimes amongst others ^5,6^. The original green fluorescent protein (GFP) from *Aequorea victoria* (avGFP) was, and still is, the underlying basic protein scaffold by which many FPs spanning the blue to yellow region are engineered ^7–10^. One of the first truly useful engineered versions of GFP still widely used today, EGFP (enhanced GFP) ^11,12^, resulted in a shift in the dominant ground state chromophore from neutral protonated form (CroOH; λ_max_ 395 nm) to the deprotonated charged phenolate form (CroO^-^) with more favourable excitation and fluorescence properties. Since then, many derivatives of avGFP have been engineered (see FPBase ^13^ www.fpbase.org/protein/avgfp/), including superfolder GFP (sfGFP) ^14^. While sfGFP folds and matures quicker, and is very stable, key fluorescence properties such as brightness and photobleaching resistance are similar to EGFP (Table S1)^15^. Brightness can be improved for green-yellow versions of avGFP (e.g. Venus^9^ and Clover^16^) and related FPs (e.g. mNeonGreen^17^) but results in reduced resistance to photobleaching (Table S1)^15^.

One of the key determinants of avGFP’s fluorescence properties is the histidine at residue 148. Together with a structured local water molecule (here termed W1), H148 directly interacts with the chromophore phenol group and is thought to play a key role in promoting and stabilising CroO^-^ in the ground state^18–24^ (Figure 1a) ^18,22,23,25^. H148 is also thought to be dynamic and has been observed to sample an alternate “open” conformation distant from the CRO ^18,21,22,26^. In many non-avGFPs, histidine is replaced with a non-ionisable polar residue^27^ (Figure S1). For example, Dronpa ^28^ has a high QY (0.85) with S142 acting as the equivalent of H148; mCherry (as with many DsRed derived FPs) also has a serine at the position equivalent to H148. The recently developed high brightness and photobleaching resistant StayGold FPs ^29,30^ have an asparagine (Asn137). Thus, H148 seems to be an uncommon choice in natural FPs for the crucial role played by this residue.

**Figure 1.**
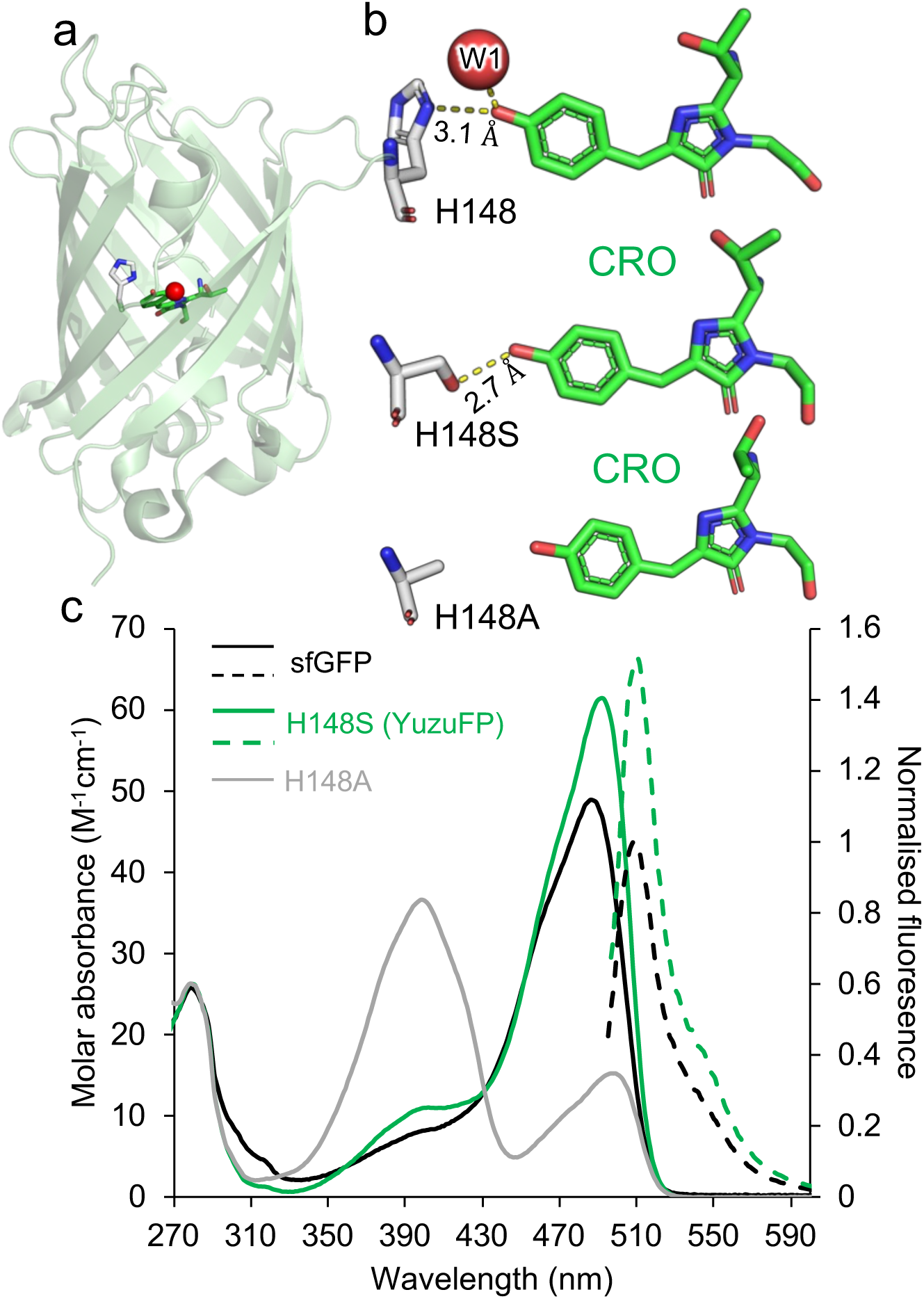
Engineering the interactions between residue 148 and the chromophore in sfGFP. (a) Overall structure of sfGFP (PDB 2b3p) ^14^ with the arrangement of the chromophore (green sticks) relative to H148 (grey sticks) shown. (b) Comparison of the H148 interaction from the crystal structure of sfGFP and the short timescale molecular dynamics models of the H148S and H148A variants. Dashed yellow lines represent polar interactions. Other H148X models can be found in Figure S3. (c) Spectral characteristics of sfGFP (black), our newly designed H148S (also known as YuzuFP) (green) and H148A (grey). Solid lines represent absorbance spectra and dashed line emission spectra. Emission spectra for sfGFP and YuzuFP were recorded on excitation at 485 nm and 492 nm, respectively. Emission spectra were normalised to the sfGFP emission maximum.

Here we assess how replacing the imidazole side chain of H148 in sfGFP modulates the charged state on the chromophore between CroOH and CroO^-^; replacement with a hydroxyl group of serine results in a brighter protein due to improved molar absorbance and QY. It also has improved resistance to photobleaching. Molecular dynamics simulations reveal the importance of more persistent H-bonding between residue 148 and the chromophore and longer water residency providing new insight into the molecular mechanism and leading to possible routes forward in how to engineer FPs for improved fluorescence properties.

## Results and discussion

### Importance of residue 148

The role played by H148 in promoting CroO^-^ in avGFP-derived proteins is well established ^18,22–24,31–33^. H148 is thought to be dynamic ^18,22,26^ capable of occupying different conformations some of which break the interaction with the chromophore ^34–37^. The most commonly observed form in crystal structures is shown in Figure 1a but the configuration is non-optimal in terms of the plane of interaction (∼140°) and distance (3.1-3.5 Å) between the imidazole side chain and chromophore (Figure S2). Even over short-time scale (10 ns) simulations the H-bond between the chromophore and H148 is present for less than 5% of the time (Table S2) and the structurally conserved nearby water molecule, W1 (Figure 1b), has a low residency time (Table S3).

The question then arises as to the importance of H148 and its imidazole side chain to avGFP fluorescence: is histidine the optimal residue? Mutant structure modelling using approaches such as AlphaFold^38^ and Rosetta^39^ is hindered for FPs by their inability to incorporate the chromophore, and crucially for FPs, the local solvation environment. Furthermore, such static modelling does not consider structural relaxation and dynamic flux. Therefore, we initially undertook molecular modelling involving short timescale molecular dynamics (10 ns) to sample all 19 residues in place of H148 (Figure 1b and S3). Modelling suggests that apart from H148, only serine and asparagine retain a side chain configuration capable of making a polar interaction with the CRO phenol group. The H148S model suggests the serine hydroxyl group forms a better configured H-bond with the CRO phenol oxygen, with a shorter distance (2.7 Å for the O-to-O distance (Figure 1b and S2) and the H-bond groups being close to planer (Figure S2). H-bond analysis also suggests the chromophore forms a H-bond with S148 more frequently and even forms two H-bonds occasionally (Table S2). In contrast H148T mutation does not appear to form a persistent H-bond via the side-chain hydroxyl. Longer distances to the phenol hydroxyl group and different relative orientations are observed for H148N (Figure S3). Starting models of H148N suggests that either the amine or carbonyl can form the polar interaction depending on carboxamide rotation. However, after 10 ns of MD only the amine forms a persistent H-bond (Figure S3 and Table S2). Replacement of H148 with alanine retains the β-carbon facing towards the chromophore but removes any polar interaction potential to the chromophore (Figure 1b and Table S2) while the larger atomic radius of sulphur results in the cysteine side-chain flipping out (Figure S3 and Tablse S2). All other mutations should remove the direct interaction with the chromophore (Figure S3).

We also considered H-bond frequency of the structurally conserved water molecule W1 (highlighted in Figure 1b) over the 10 ns simulation. In general, as with the sfGFP WT, W1 quickly (within ∼2 ns) diffuses away from its original position (Table S3). With regards to sfGFP, W1 is H-bonded to the chromophore for just under 6% of time (Table S3). In comparison, W1 H-bonds to the chromophore more frequently in the H148S (19.6%), H148A (22.6%) and H148T (13.1%) mutant simulations. Both forms of H148N together with H148C model simulations show similar H-bond occurrence as sfGFP WT (Table S3). Therefore, the short timescale MD modelling data suggests that the H148S mutation has the highest propensity to populate local H-bonds with the chromophore phenol group.

### The effect of mutating H148 on the spectral properties of sfGFP

Based on our modelling, we generated H148A, H148C, H148N, H148S and H148T mutants of sfGFP to confirm how our simulation-based modelling approach can help predict the impact of mutations on polar interactions with the CRO. The H148T mutant in our hands did not generate any fluorescent protein suggesting the mutation impacted folding and/or chromophore maturation. H148A and H148C results in a mixed population comprising a dominant CroOH peak (∼400 nm) and minor CroO^-^ peak (∼490 nm) (Figures 1b and S4a and Table S4). This confirms that removing the potential to form a polar bond between residue 148 and the CRO shifts the majority chromophore form sampled to CroOH. It was thought that residue 65 comprising the chromophore was the prime driver of chromophore ground state charge^40^. All variants emit at ∼510 nm irrespective of excitation at either ∼400 nm or ∼490 nm (Figure S4b-d and Table S4), with excited state proton transfer the likely mechanism for the large Stokes shift of the CroOH ground state starting point^23^. The H148N sfGFP mutant had similar absorbance spectrum to WT sfGFP (Figure S5) so retaining the CroO^-^ ground state, as suggested would be the case in the modelling. Previous work had suggested that H148N should promote the CroOH despite being able to make H-bond contact with the phenol hydroxyl group ^41^. The overall spectral properties of sfGFP-H148N are no better than the parental sfGFP (Table S4) so was not explored further.

The H148S mutation resulted in a variant that had higher molar absorbance and Quantum Yield (QY) compared to sfGFP, with the variant being 1.5 fold brighter than sfGFP itself (Figure 1c, Table 1) and 1.7 times brighter than EGFP. The sfGFP H148S protein, termed YuzuFP from here on in, has a slightly red shifted excitation and emission (Table 1) giving the protein a yellow hue compared to sfGFP, similar to the Yuzu fruit. YuzuFP has a Stokes shift of 19 nm, which is larger than the recently developed StayGold^42–44^ variants (7-10 nm), meaning there is less spectral overlap between the peak excitation and emission events. The fluorescence lifetime of YuzuFP (2.76ns) is also slightly longer than both sfGFP (2.51 ns; Table 1 and Figure S6a) and EGFP (2.6 ns). Mutation of H148 does not significantly effect the pKa of YuzuFP (6.2), which is similar to sfGFP and EGFP (Table 1 and Figure S6b). This suggest that H148 is not playing a major role in dictating pH dependent avGFP fluorescence. YuzuFP fluorescence was not adversely affected by salt (NaCl or KCl; Figure S6c-f) and is also largely monomeric in line with the parental sfGFP (Figure S7).

**Table 1.**
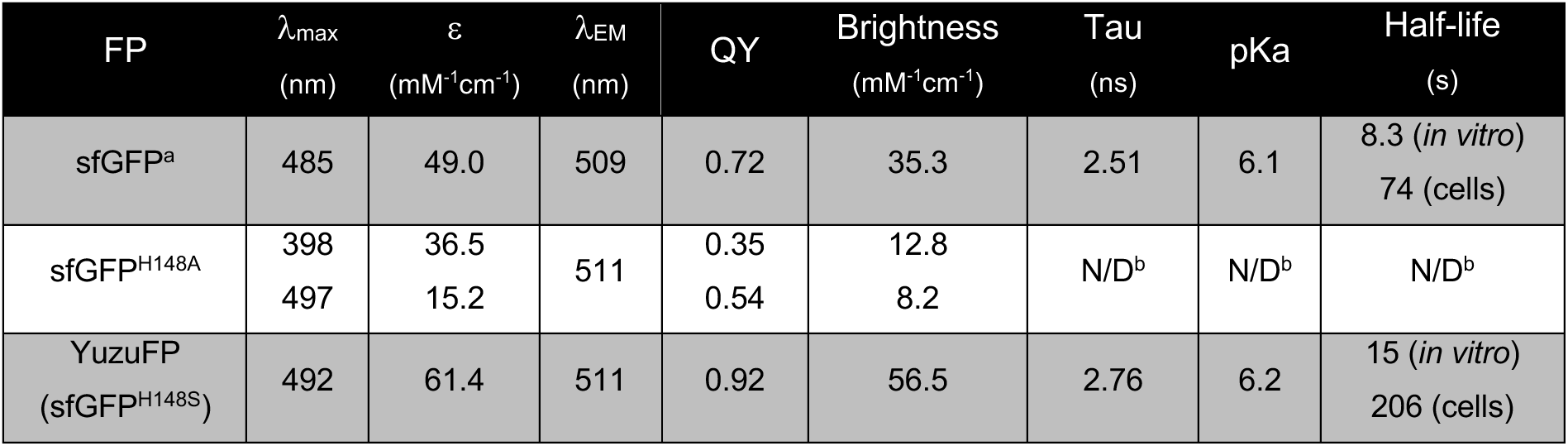
Spectral properties of sfGFP-H148X variants. a, originally reported by Reddington et al. ^31^ b, N/D denotes not determined.

| FP | $\lambda_{\text{max}}$<br>(nm) | $\epsilon$<br>(mM<sup>-1</sup>cm<sup>-1</sup>) | $\lambda_{\text{EM}}$<br>(nm) | QY | Brightness<br>(mM<sup>-1</sup>cm<sup>-1</sup>) | Tau<br>(ns) | pKa | Half-life<br>(s) |
| --- | --- | --- | --- | --- | --- | --- | --- | --- |
| sfGFP<sup>a</sup> | 485 | 49.0 | 509 | 0.72 | 35.3 | 2.51 | 6.1 | 8.3 (*in vitro*)<br>74 (cells) |
| sfGFP<sup>H148A</sup> | 398 | 36.5 | 511 | 0.35 | 12.8 | N/D<sup>b</sup> | N/D<sup>b</sup> | N/D<sup>b</sup> |
|  | 497 | 15.2 |  | 0.54 | 8.2 |  |  |  |
| YuzuFP<br>(sfGFP<sup>H148S</sup>) | 492 | 61.4 | 511 | 0.92 | 56.5 | 2.76 | 6.2 | 15 (*in vitro*)<br>206 (cells) |

As there is a shift towards super-resolution single molecule imaging which generally requires higher power inputs, we next undertook single FP analysis of sfGFP and YuzuFP using TIRF microscopy^45^. YuzuFP has improved single molecule characteristics compared to sfGFP; comparison of individual traces shows that overall YuzuFP remains fluorescent over longer timescales than sfGFP (Figure 2a and Figure S8). This equates to YuzuFP being more resistant to photobleaching (Figure 2b) with an ensemble half-life under our single molecule imaging conditions of 15.0 ± 0.1 s, ∼2 fold longer than WT sfGFP (half-life 8.3 ± 0.1 s). In line with the increased half-life, the lifetime increases from 12.0 ± 0.2 s for sfGFP to 21.5 ± 0.1 s. Analysis of individual traces show that a significant number of YuzuFP molecules (24%) remained fluorescent for >25 s (Figure 2a and S8) at sufficient laser powers for dynamic single molecule imaging at a time resolution of 60 ms (see Methods). As sfGFP is already one of the most photostable green-yellow FPs ^15^, YuzuFP represents a marked improvement (see Table S1 for examples).

**Figure 2.**
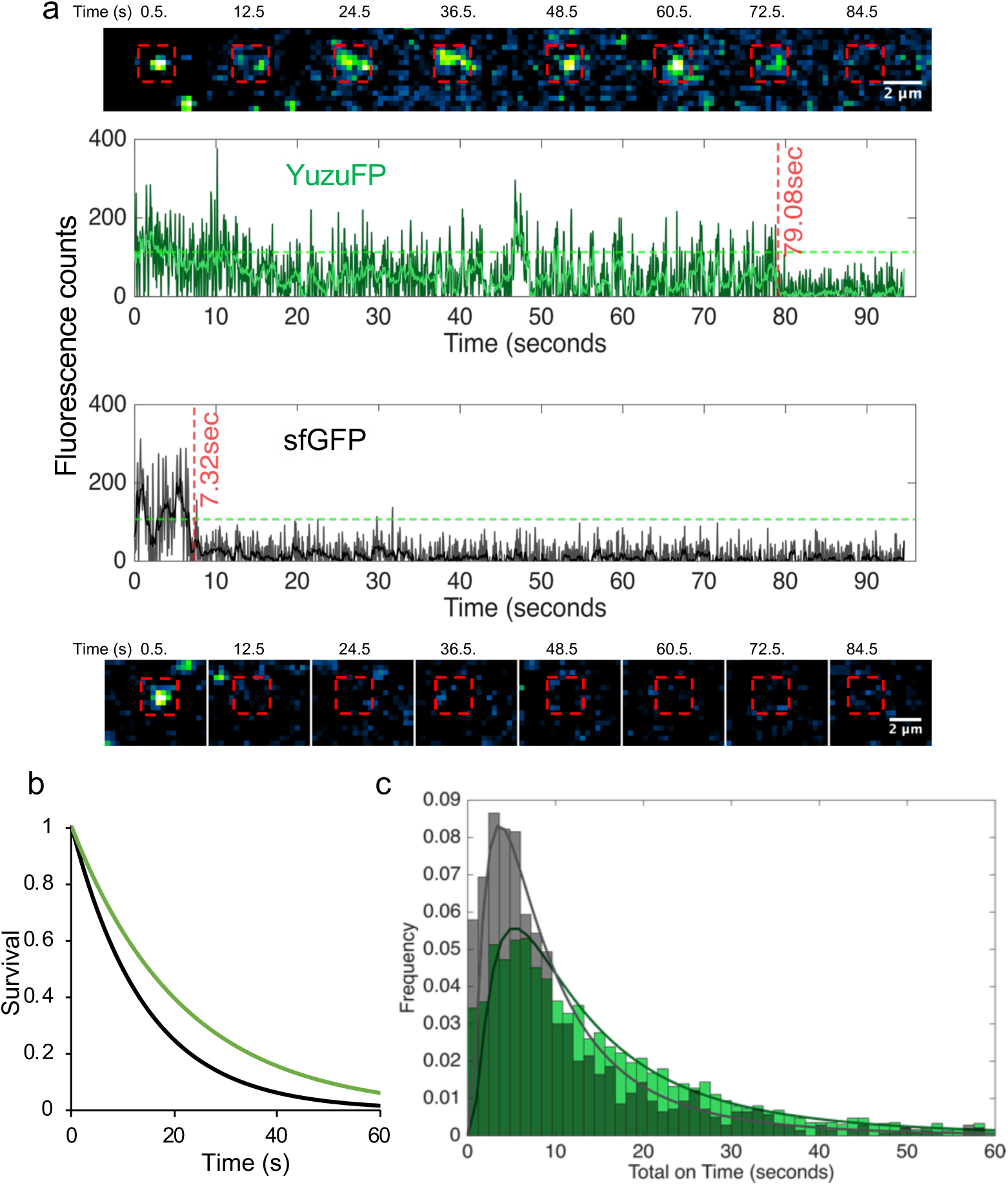
Single molecule fluorescence characteristics of sfGFP and YuzuFP. (a) Example single FP fluorescence images at various timepoints and their corresponding emission traces for YuzuFP (top, green) and sfGFP (bottom, black). In the image time course, the red box denotes the region of interest representing the analysed area. Dark green and grey traces show the raw data for each protein while the lighter green and black lines show the result of a forward-backward moving window filter (Chung-Kennedy) applied to the raw data. The green dashed line in each instance represents the threshold separating values considered on and off. More traces can be found in Figure S8. (b) Single molecule photobleaching survival plot for YuzuFP (green) and sfGFP (black) based on proportion of FPs retaining fluorescence (survival) over time. The decay curves were generated by fitting a single component exponential function to empirical cumulative distribution functions comprised of lifetimes from 3766 and 1283 individual FP traces for Yuzu FP and sfGFP, respectively. (c) Frequency distribution of single molecule photobleaching lifetimes for YuzuFP (green) and sfGFP (grey), representing the time spent in the on-state prior to photobleaching. The data was binned at 1.2 second intervals. The solid lines represent a log-normal fit to the distribution.

FPs observed at the single molecule level have a propensity to “blink” switching between on and off states. Previously the presence and type of this blinking has been indicated as a contributing factor in the manifestation of ensemble level behaviours. YuzuFP is shown to cumulatively spend a longer time in its on state than sfGFP with the mean on-time peak shifting from 3.6 s to 5.3 s (Figure 2c).

To confirm that *in vitro* performance translates to *in situ* cell imaging, a Lifeact-YuzuFP fusion was imaged by wide-field fluorescence microscopy in live cells. HeLa cells expressing the LifeAct fusions clearly show actin filament structures as expected (Figure 3a-b). While the ability to resolve filament structure is lost at 40 s for sfGFP-LifeAct fusions, YuzuFP-LifeAct retains the ability to visualise clear filaments structures up to 120 s in line with the improved photostability. The photobleaching half-life is calculated to be nearly 3-fold higher for the YuzuFP fusion compared to sfGFP (206 ± 3.7 s *versus* 74 ± 0.5 s, respectively) with corresponding increases in lifetimes (106 ± 0.7 s for sfGFP and 297 ± 5.2 s for YuzuFP) confirming YuzuFP’s superior photostability over the already stable sfGFP starting point.

**Figure 3.**
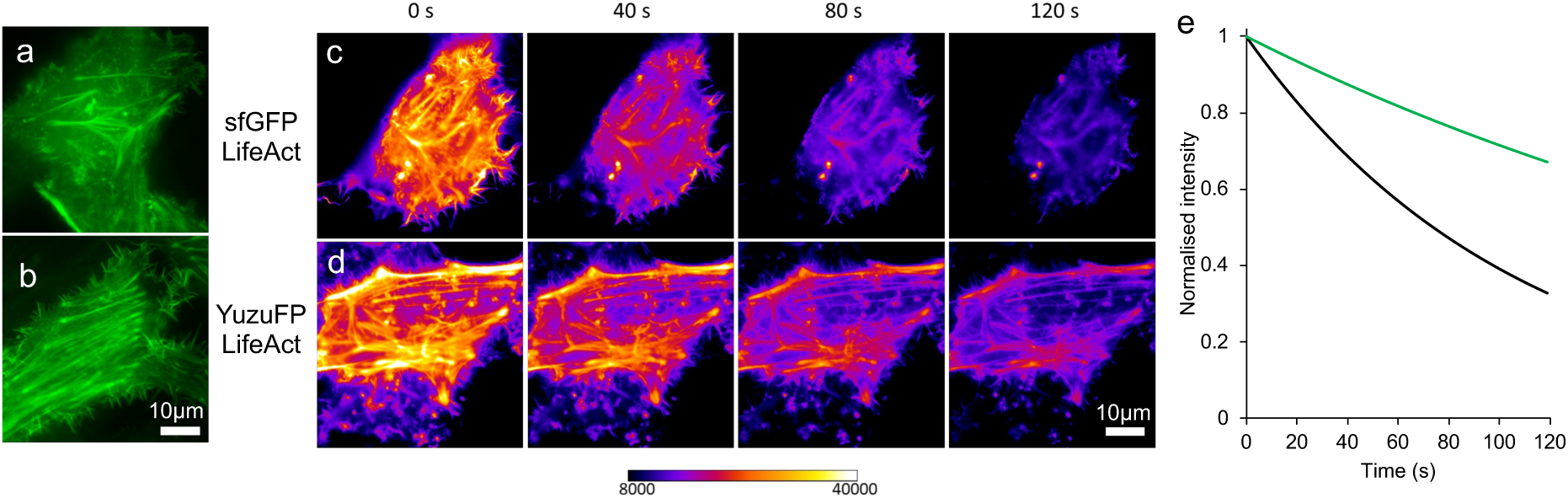
Live cell imaging of YuzuFP and sfGFP LifeAct Fusions. Widefield image of (a) sfGFP and (b) YuzuFP LifeAct fusions. Time course of (c) sfGFP and (d) YuzuFP LifeAct fusions false coloured using a 16 bit Fire look-up table with the black value set to 8000, and the white value set to 40000, to visualise signal within the dynamic range of the microscope setup. Note the similar intensities following 40s constant exposure of sfGFP, to the 120s timepoint of YuzuFP. (e) *In situ* photobleaching of YuzuFP (green line) and sfGFP (black line). Six independent intensity values were normalised to 1.0 (based on the intensity at 0 s), background subtracted and fit to a one phase decay curve in GraphPad Prism.

### Long scale molecular dynamics reveals potential beneficial effect of H148S mutation

To investigate how the H148S mutation could be exerting its influence, we extended the initial MD simulations to 3 repeats of 500 ns on the YuzuFP CroO^-^ model and compared to sfGFP. Comparison of root mean square fluctuation (RMSF) shows both proteins display relatively similar backbone fluctuations (Figure S9a), with most differences being on the sub Ångstrom level (Figure 4a). The chromophore does not undergo any major structural change over the course of the simulation for both sfGFP and YuzuFP (Figure S9b-c). The root mean square fluctuation (RMSF) of the hydroxyethyl group of the original Thr65 together with the original backbone oxygen of Gly67 show some fluctuation but most CRO atoms remain relatively unperturbed (Figure S9d).

**Figure 4.**
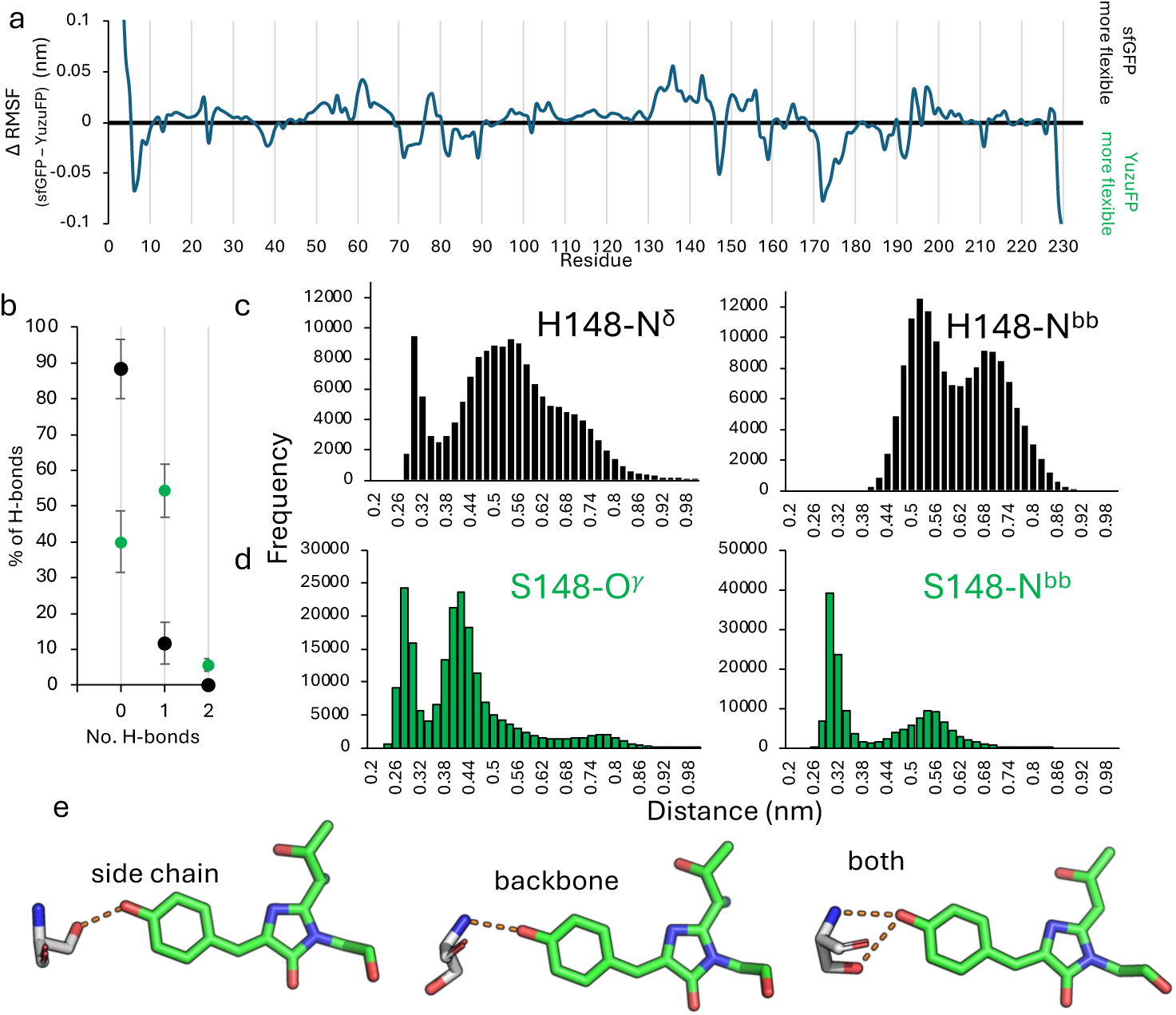
Hydrogen bonding between the chromophore and residue 148. (a) Per residue root mean square fluctuation (RMSF) Cα difference plot. The difference plot was generated by subtracting YuzuFP from sfGFP. The individual RMSF plots are shown in Figure S9a. Positive values indicate that sfGFP is more flexible and negative values that YuzuFP is more flexible. (b) The percentage of time H-bonds are formed between CRO and residue 148 over the course of the MD simulations for sfGFP (black) and YuzuFP (green). Error bars are the standard deviation between values measured for the 3 individual simulations. The pair-wise distance distribution in (c) sfGFP and (d) YuzuFP, between chromophore phenol oxygen and residue 148 backbone and side chain H-bond donor heavy atoms across all simulation data. Bin sizes are 0.02 nm. The distance against time plot is shown in Figure S10. (e) Representative individual trajectories illustrating the H-bond between the S148 sidechain hydroxyl group (simulation 2, 297.80 ns), backbone amine group (simulation 2, 254.39 ns) or both in YuzuFP (simulation 2, 5.43 ns). H-bonds are shown as orange dashes.

One major difference is the interaction between CRO and residue 148. With regards to sfGFP, H-bonding occurs less frequently than with S148 in YuzuFP. H148 does not make H-bond contact with the chromophore for *circa* 88% of the simulation time while in YuzuFP the S148 H-bonds with CRO for *circa* 60% of the simulation time, with ∼5.5% comprising of two H-bonds (Figure 4b). This is similar to that observed over the initial 10ns simulations (Table S2). The change in H-bond frequency is manifested in the pairwise distance distribution between the CRO phenol O atom and either the side chain (Nδ atom in sfGFP H148 or Oγ atom in YuzoFP S148) or the backbone amide N (Figure 4b-c). The sidechain of residue 148’s H-bond group is on average 1 Å closer to the CRO phenol oxygen in YuzoFP than sfGFP (4.2 ±1.3 *versus* 5.2 ±1.4). It is also clear from the atom pair distance distributions that the backbone amide N of S148 in YuzoFP is more consistently within H-bond distance of the CRO phenol oxygen (<3.5 Å) while H148 in sfGFP rarely comes within 4 Å (Figure 4c-d); the average atom pair distance between the amide nitrogen and chromophore phenol oxygen being 4.0 ±1.2 Å in YuzoFP compared to 6.1 ±1.0 for sfGFP. The backbone amide group of S148 in YuzuFP can form a H-bond with the chromophore phenol oxygen, which is not observed for sfGFP. Indeed, on average just over 50% (55% ±21) of H-bonds between S148 and involve the backbone amide. This suggests that the H148S mutation allows switching between the side chain and backbone H-bonding with the chromophore, and on occasion both for ∼5% of the time. This is clearly seen from individual trajectories whereby either the side chain, backbone or both are making polar contacts with the CRO phenol oxygen (Figure 4e).

The reason for the change in H-bond interactions appears to be largely down to changes in orientation and distance of residue 148 with respect to the chromophore. The root mean square deviation (RMSD) for H148 in sfGFP for most of the simulations appears to be relatively stable but periodically undergoes a significant change (> 0.1 nm; Figure 5a). The RMSD of S148 in YuzuFP quickly converts to a form with an RMSD of *circa* 0.8 nm and fluxes periodically with a conformation closer to that of the start point (Figure 5b). Distances between the chromophore phenol oxygen and the Cα of residue 148 suggest that in YuzuFP, the change in RMSD can be ascribed to S148 moving closer to the chromophore while H148 in sfGFP moves further away (Figures 5c); the average distance between the 148 Cα and chromophore phenol oxygen is 4.45 Å ± 0.80 for YuzuFP, circa 1.5 Å shorter than in sfGFP (5.93 Å ±0.87). The higher RMSD observed for YuzuFP does not impact S148’s ability to make H-bonds with the chromophore (Figure 5d). The larger periodic deviations observed for H148 in sfGFP mirrors the alternative “open” conformation observed in crystal structures (Figure 5e) ^34–37^. Thus, the closer association of S148 with the chromophore together with a high propensity to form H-bonds is likely to play a key role in improving the properties of YuzuFP.

**Figure 5.**
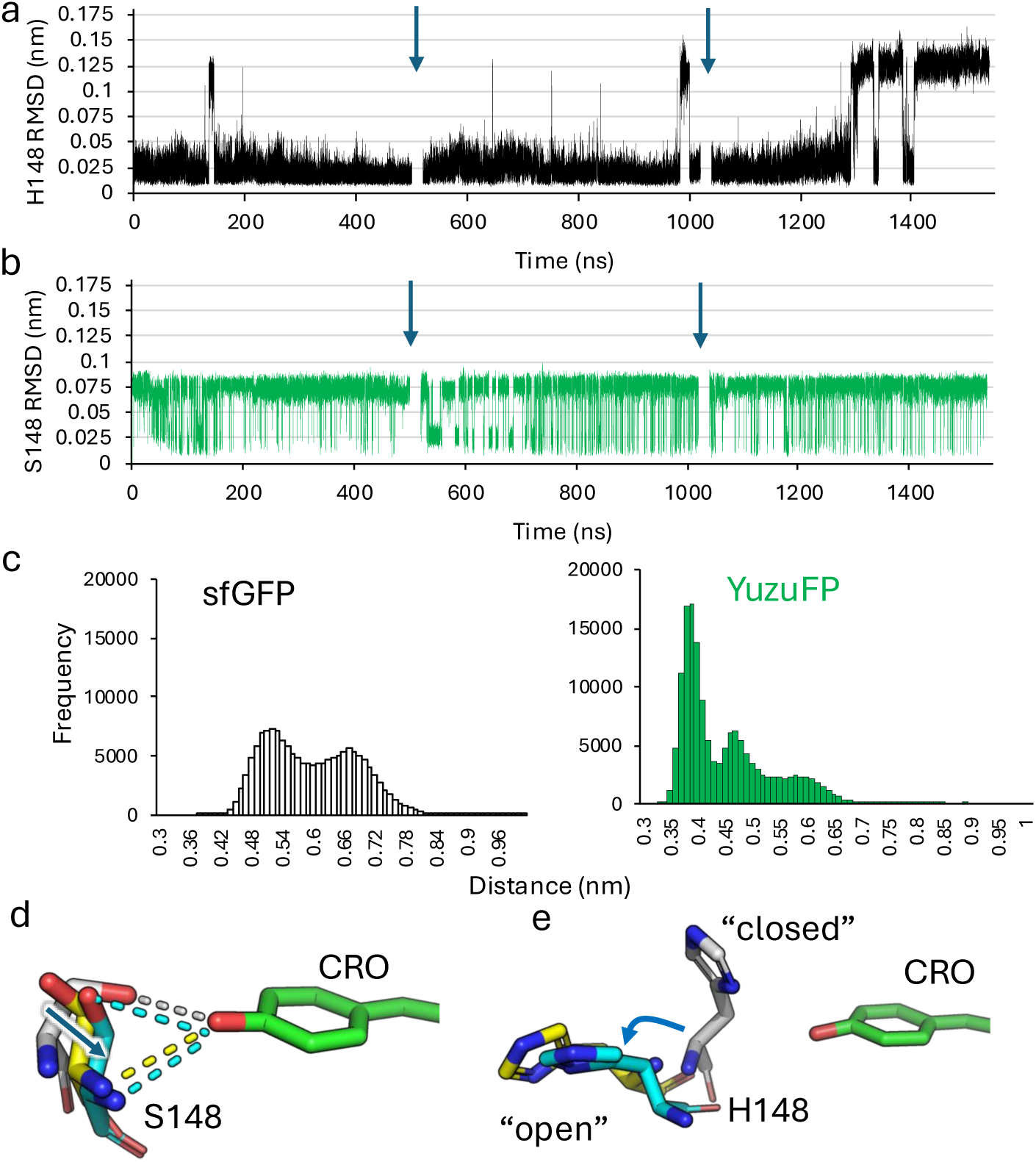
Conformational changes to residue 148. The root mean square deviation (RMSD) of (a) H148 in sfGFP and (b) S148 in YuzuFP. (c) The distance distribution between the CRO phenol oxygen and the Cα of residue 148 for all simulations. Bin size is 0.01 nm. The distance change over time plot for each simulation is shown in Figure S11. (d) Representative trajectories of YuzoFP (grey, 0 ns; yellow, simulation 1 150.75 ns, RMSD 0.0894 nm; cyan, simulation 1 168.74 ns, RMSD 0.08 nm). The dashed lines represent polar interactions. (e) Representative trajectories of sfGFP (grey, 0ns; yellow simulation 3, 262.86 ns, RMSD 0.12 nm; cyan, simulation 3, 250.87 ns RMSD 0.12 nm). The blue arrow signifies direction of backbone movement with respect to the starting trajectory (0 ns).

### Dynamic water molecules

The chromophore while being internal to the barrel structure is well solvated with sfGFP having six water molecules surrounding the chromophore in the original crystal structure of sfGFP ^14^ (Figure S12). One water molecule, termed W1 in Figure 1a, is ever present in crystal structures of avGFPs (Figure 1b) and other FPs (Figure S1) forming polar interactions with CRO phenol O as well as the side chain of S205; it’s positioning is thought to be important in promoting the CRO-O^-^ ^32,46^ state through proton shuttling and enhancing fluorescence ^23,47–49^. Our MD simulations show that W1 is associated with the protein for a relatively short time at that spatial position in both sfGFP and YuzuFP before exchanging with bulk solvent (Figure 6). The residency time of W1 for YuzoFP is longer (*circa* 8.5 ns) compared to sfGFP (*circa* 1.5ns). The analysis is complicated as in one sfGFP simulation, W1 quickly becomes internalised (within 0.22 ns) close to the CRO (Figures 6b and S13). W1 does appears to be replaced through the simulation with other water molecules; the CRO phenol oxygen in YuzuFP spends only 5.7% of its time with no H-bonds to water, which increases slightly to 6.8% in sfGFP.

**Figure 6.**
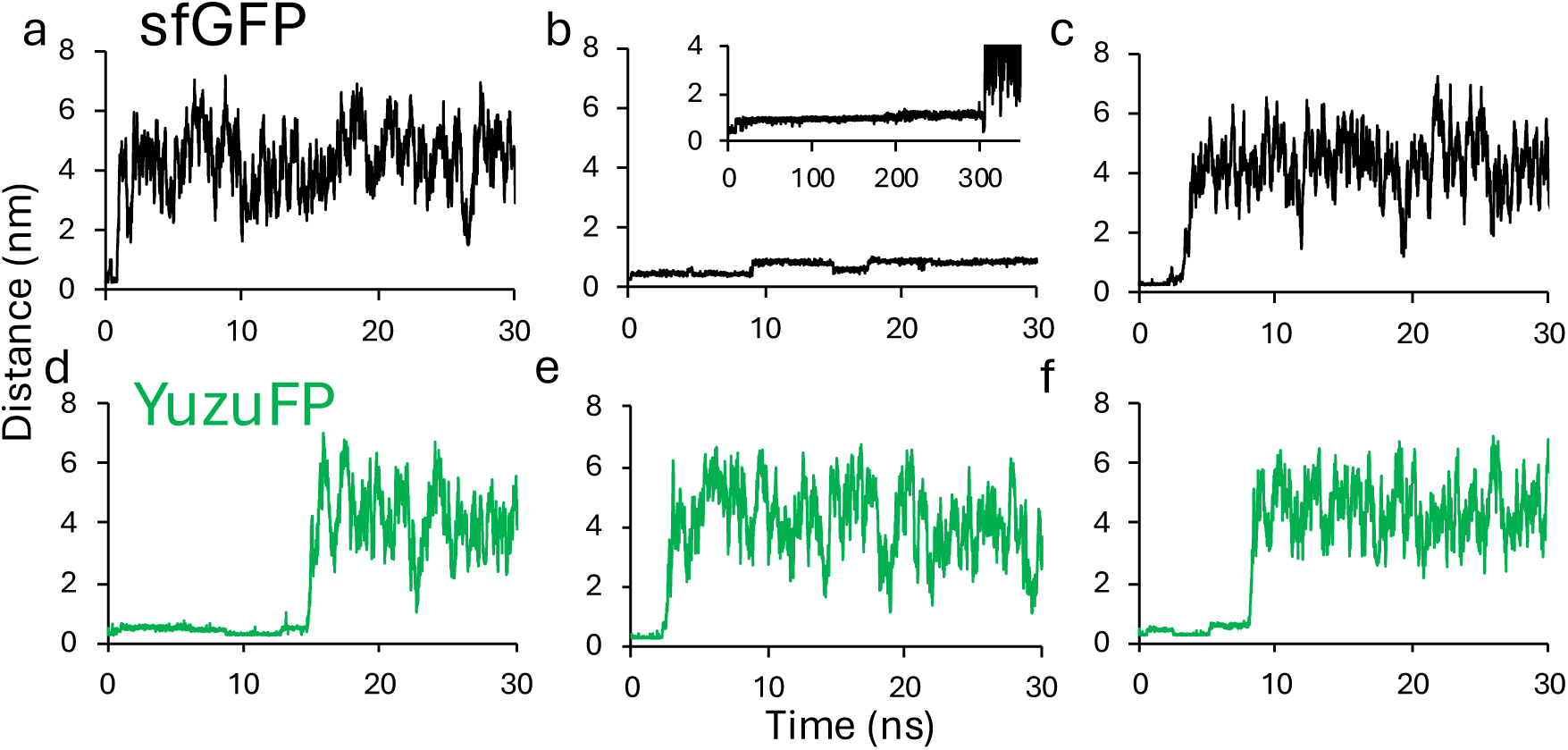
Residency of structurally conserved water molecule W1 close to chromophore phenol oxygen. Distance between chromophore phenol oxygen and O atom of water molecule W1 (see Figure 1a for reference) in (a-c) sfGFP and (d-e) YuzoFP in each of the 500 ns simulations. In 6b, shown inset is the distance over an extended period (350 ns).

The remaining water molecules that reside within the barrel structure overall have longer residency times but most eventually undergo bulk exchange with the solvent (Figure S14). Analysis of potential tunnels using CAVER ^50,51^ reveals an exit point close to H148 in sfGFP (Figure S15) providing a potential route to exchange with bulk water. However, some of these water molecules reside close to the CRO over the course of simulation. For YuzoFP water W2 remains buried in two simulations whereas it exchanges within 172 ns in sfGFP (Figure S14 and Table S5). In the third YuzoFP simulation two other water molecules (W5 and W6) remain buried over the 500 ns (Figure S14 and Table S5). In contrast, only two waters in a single simulation remain buried over the simulation for sfGFP (Figure S14 and Table S5). Water exchange rates with the bulk solvent could be indicative of chromophore accessibility to other solutes such as reactive oxygen species that cause photobleaching. Thus, mutations to residue 148 that affect local dynamics could also influence tunnel aperture and thus access to the chromophore.

## Conclusion

The avGFP still provides the basic framework by which many FPs are engineered but H148 remains largely a constant in green-yellow versions despite FPs from other origins using different residue types in the same capacity. Using a molecular dynamics based approach we were able to model the potential changes to chromophore interactions on mutating H148 to each other amino acid in water environment, something not achievable by classical protein design approaches. Short scale molecular dynamics suggested H148S mutation will generate a more persistent H-bond with the chromophore than the native histidine. By mutating H148 to serine in the sfGFP background, the new engineered FP termed YuzuFP is brighter, more resistant to photobleaching, and stays on for longer making it a good candidate for modern fluorescence microscopy approaches. The origin of the improvement appears to be changes in the local dynamics around residue 148 with mutation to serine resulting in a more stable and persistent interaction with the chromophore and water molecules. Thus, YuzuFP could prove to be an important tool for researchers and together with MD-based design could form the basis for future protein engineering endeavours aimed at further improving its utility such as brightness.

## Methods

### Protein engineering and recombinant production

The sfGFP variants in the pBAD plasmid were generated using a whole plasmid, inverse PCR process as described previously ^31,32^ using the primers in Table S6 and the Q5^®^ site-directed mutagenesis kit from New England Biolabs. Recombinant production was performed in *E. coli* essentially as described previously ^31^. Chemically competent *E. coli* TOP10™ cells were transformed with the prerequisite plasmid and plated on LB agar plates supplemented with 50 μg/ml ampicillin. A single colony was taken from LB agar plate and used to inoculate a 10 mL 2xYT starter culture supplemented with and incubated overnight with shaking. The overnight culture was used to inoculate a 1L culture of autoinduction media (see supporting methods). The culture was inoculated overnight at 37 °C with shaking. Cells were then pelleted by centrifugation and suspended in 50 mM Tris, pH 8.0. Cells were lysed using the French Pressure Cell. Soluble cell lysate was then separated from insoluble fractions via centrifugation using a Beckman JLA 25.50 rotor at 25,000 x g for 40 mins. Protein purification was carried out with an ÄKTA Purifier FPLC using columns purchased from Cytiva and protein elution monitored at 280 nm, 400 nm and/or 485 nm. Clarified cell lysate was passed through a 5 mL His Trap^TM^ HP column (binding capacity ∼200 mg protein) equilibrated in 50 mM Tris, pH 8.0 buffer containing 10 mM imidazole. Bound target protein was then eluted by the addition of the elution buffer containing imidazole at a gradient from 10 to 500 mM imidazole. Fractions were checked for purity via SDS-PAGE analysis. Finally, the proteins were further purified by size exclusion chromatography (SEC) using a HiLoad^TM^ 16/600 Superdex^TM^ S75 pg column (Cytiva) equilibrated with 50 mM Tris, pH 8.0. Protein samples were concentrated using Vivaspin^TM^ 10 kDa molecular weight cut-off spin filters (VWR) by centrifugation at 4000 x g until the desired volume is reached. Analytical SEC was performed using a Superdex 75 10/300GL column (Cytiva).

### Spectroscopic analysis

Protein concentrations were determined using the Bio-Rad DC Protein Assay using sfGFP as the standard. Concentration was confirmed by comparison to sfGFP’s 280 nm absorbance (ε_280_ = 25.3 mM^-1^cm^-1^); as the mutants did not contain any additional UV absorbing side chains their 280 nm molar absorbance should be the same as for sfGFP (see Figure 1c). UV-visible (UV-vis) absorption spectra were recorded on an Agilent Cary 600 spectrophotometer in a 1 cm pathlength cuvette. Spectra was recorded from 200 – 600 nm at a rate of 300 nm/min using a known protein concentration (usually 5 μM) and the Beer-Lambert equation used to determine the molar absorbance of each variant protein. Emission and excitation spectra were measured using a Varian Cary Eclipse Fluorimeter with a QS quartz cuvette (Hellma). Data were collected with 5 nm slit width at a rate of 300 nm/min. Emission spectra were recorded at a fixed excitation wavelength according to the excitation maximum of the variant. Spectra were recorded using 0.5 μM of protein in 50 mM Tris-HCl, pH 8.0. Quantum yield was determined as described previously ^24^ using a fluorescein standard (QY = 0.75 in 0.1 M NaOH). The pKa was determined using changes in absorbance at the λ_max_ equivalent to the CroOH (∼400 nm) and CroO^-^ (∼490 nm) using the following buffers: 100mM Glycine-HCl (pH 3.0), 100mM acetate buffer (pH 4.5 to 5.5), 100mM KH_2_PO_4_-NaOH (pH 6.0), 100mM Hepes buffer (pH 7.0 and 8.0), 50 mM TrisH-Cl (pH 8.0), 100mM Glycine-NaOH (pH 9.0-10.0), 100mM Na_2_HPO_4_-NaOH (pH 11-12). Fluorescence lifetime measurements were performed using a bespoke device setup described previously ^52^ at protein concentrations ranging from 5-15 μM. The resulting decay curves were fit to a single exponential decay using GraphPad Prism.

### Single molecule analysis

Single molecule imaging of YuzuFP and sfGFP were carried out similar to that as described previously ^32^. Briefly, total internal reflection fluorescence (TIRF) imaging of a low concentration FP solution on a plasma treated coverslip was performed using a custom optical setup. The core of this is a Nikon Ti-U inverted microscope with a high numerical aperture 60x objective (Nikon, CFI Apochromat TIRF) and an Andor iXon ultra 897 camera. Excitation was achieved using a fibre-coupled Venus 473nm DPSS laser with a power output of 100mW (achieving ∼7.8μW/μm^2^ at sample). A 488nm dichroic mirror coupled with a 500nm long pass filter and a 525/50nm bandpass filter were used to isolate the fluorescence signal. Acquisitions were made in areas without prior laser exposure to minimise the effects of photobleaching prior to image capture. An exposure time of 0.06 seconds and an acquisition period of 96 seconds (1600 frames) was used in each experiment. Acquired image stacks were processed using the FIJI distribution of ImageJ to normalise for laser power fluctuation and spatial variations in laser intensity as previously described ^32,53^. Detection and extraction of single molecule time series data was also carried out in FIJI using the TrackMate plugin^54,55^ employing a difference of gaussians detection method and an estimated spot size of 4 pixels. Extracted time series were exported to MATLAB and a forwards backwards moving window Chung Kennedy filter^56^ was applied. An intensity threshold for each time series was determined by first segmenting using a minimum sum log-likelihood deviation from segmented means method^57^. Values from the lowest intensity segment determined by this method were averaged and an intensity threshold was determined as 9 standard deviations from the mean. A second round of temporal thresholding was applied to values meeting the intensity threshold criteria. A window size of 15 frames (0.9 seconds) was used as a lower limit to separate on-state events from background fluctuations in noise. Intensity values showing sequential temporal separation below this threshold were grouped into single events. Photobleaching lifetime was determined as the end point of the last of these events in a given time series. Cumulative on-time was taken as the sum of all on-events within a given molecule’s photobleaching lifetime. Mean on-time of the two species was determined by fitting a log-normal distribution to a probability density function generated from all on-times from each FP using GraphPad Prism. Similarly, the survival half-life was determined by fitting a single component exponential decay to the empirical cumulative distribution function of photobleaching times for each FP. Datasets for the two FPs were made up of three experimental repeats each and a total of 3766 and 1283 single molecule spots and corresponding time series were extracted from the YuzuFP and sfGFP datasets respectively.

### Cell imaging

The LifeAct-sfGFP fusion gene was synthesised by Twist Biosciences (see Supporting Information for sequence) and placed in their pTWIST CMV mammalian expression vector. Human cervical carcinoma (HeLa) cells (ATCC, UK) were cultured directly onto live cell dishes (Mattek, USA), and allowed to adhere overnight before transfecting using Fugene6 (Promega) according to manufacturer’s instruction using 1 μg of DNA per 3 μl Fugene in a volume of 100μl of OptiMEM (Life Technologies, UK). Before imaging, cells were washed and the media replaced with Fluorobrite DMEM containing 10% FBS (Life Technologies, UK). Wide-field epi fluorescence measurements were conducted on an inverted Olympus IX73 microscope and a Prior Lumen200Pro light source using filter set (89000, Chroma, Vermont, U.S.A.) selecting the ET490 nm/20 nm as the excitation filter and the ET525 nm/36m nm as the emission filter. The fluorescence emission was detected with a Hamamatsu ORCA-flash 4.0 V2 sCMOS Camera operated utilizing the HCImage software package (Hamamatsu). A 100× oil immersion objective with an NA of 1.4 was used to collect sequential images of 1 s exposure over 121 timepoints. Cells were imaged live at 37 °C 5% CO_2_ in a humidified chamber. For quantification of intensity and photobleaching, a 10 μm diameter circular region of interest was measured for average intensity within individual transfected cells utilizing the multi measure function within the ROI manager of FIJI ^53^. All regions of interest were checked to ensure that all pixels were within the dynamic range of the microscope setup. All images were captured and processed using the same methodology. >20 ROIs per condition were assessed. Fluorescence intensity measurements were normalized to their brightest value (at timepoint 0), background adjusted and fit to single phase decay model in GraphPad Prism.

### Modelling and molecular dynamics

For the short scale MD, the initial models of every possible amino acid at position 148 were generated using the PyMOL ^58^ mutagenesis tool using the sfGFP crystal structure (PDB 2b3p) as the starting point. Short timescale molecular dynamics were performed using GROMACS^59,60^ with a AMBER99SB forcefield ^61^ modified to contain constraints for the GFP chromophore (see https://github.com/drdafyddj/GFP). The protein was then placed centrally in a cubic box at least 1 nm from the box edge. The protein system was then solvated with water molecules and total charge balanced to zero with Na^+^ ions. The protein was then energy minimized to below 1000 kJ mol^−1^ nm^−1^ with an energy step size of 0.01 over a maximum of 50000 steps. The system was then temperature and pressure equilibrated before MD runs at 300 K, 1 atmosphere pressure for 10 ns with a 2fs time step integration. Clustered average models sampling the duration of the simulations were used as represented structures of the mutations. Long scale molecular dynamics of sfGFP and YuzuFP were undertaken in the same manner but extended to 3x500 ns simulations. The original crystallographic waters present in the 2b3p PDB file were included in the long run simulations. All simulations were performed on the Super Computing Wales Hawk facility (project code scw1631). The trajectories files were centred and dumps of individual trajectories were performed via the *trjconv* command. RMSD and RMSF calculations were performed using the *rms* and *rmsf* commands. Pairwise distances and hydrogen bonds were determined using the *pairdist* and *hbond* commands.

## Code availability

Molecular dynamics was performed using the open source software Gromacs (https://www.gromacs.org). Additional parameter files are available via GitHub (https://github.com/drdafyddj/GFP).

## Acknowledgements

D.D.J would like to thank the following funders: KESS2 (Knowledge Economy Skills Scholarship European Regional Development Fund via Welsh Government) studentship in partnership with Tenovus (for R.D.A), a Royal Thai Embassy studentship to D.V., BBSRC (International Partnership Award and BB/Z514913/1), the EPSRC (EP/V048147/1). The authors would like to thank the Cardiff School of Biosciences Protein Technology Hub for helping with the production and analysis of proteins, and the Cardiff University Bioimaging Hub Core Facility for their support and assistance in this work. Molecular dynamic simulations were in run on the Advanced Research Computing at Cardiff (ARCCA) facility (Hawk cluster) as part of Supercomputing Wales part-funded by the European Regional Development Fund via the Welsh Government under project code scw1631. O.C. would like to acknowledge support for the work in part from the European Union’s Horizon 2020 research and innovation programme under grant agreement No. 824060 (project ACDC), Horizon Europe European Innovation Council Pathfinder programme under grant agreement No. 101130747 (project Bio-HOST) Co-funded by the European Union and UKRI and BBSRC project BB/Y008537/1 (ALMOND).

## Author contributions

R.D.A., W.D.J, D.V. and A.Z contributed equally to this work. The order is based on the alphabetical order of their surnames. All authors contributed to the writing of the paper and analysing data. R.D.A. contributed to the conception of the project, undertook initial modelling, produced mutants, produced protein and contributed to the steady-state spectral analysis. W.D.J. undertook single molecule measurements and analysis. D.V. contributed to mutant production, steady-state spectral analysis and long-scale molecular dynamic analysis. A.Z. contributed to mutant production, steady-state spectral analysis and mammalian cell imaging work. K.A.P contributed to the initial short scale molecular dynamics. O.K.C contributed to single molecule measurements and analysis and directed the project. P.D.W. contributed to mammalian cell imaging work and analysis, and directed the project. D.D.J. conceived and directed the project, contributed to general data analysis and long scale molecular dynamics analysis.

## Supplementary Information

### Supporting Methods

#### Autoinduction media

The following lists the auto-induction media used to produce sfGFP proteins present in the pBAD plasmid: 1% (w/v) tryptone; 0.5 %(w/v) yeast extract; 0.5% (v/v) glycerol; 0.05 % (w/v) glucose; 0.2% lactose; 25 mM Na_2_HPO_4_; 25 mM KH_2_PO_4_; 50 mM NH_4_Cl; 5 mM NaSO_4_; 2 mM MgSO_4_; 1 x trace metals (4 μM CaCl_2_; 2 μM, MnCl_2_, 2 μM ZnSO_4_, 0.4 μM CoCl_2_, 0.4 μM CuCl_2_, 0.4 NiCl_2_, 0.4 uM Na_2_MoO_4_, 0.4 μM H_3_BO_3_ and 10 μM FeCl_3_ in ultra-pure water) and 0.05 % (w/v) L-arabinose. The medium was supplemented with 50 μg/ml ampicillin. All chemicals were supplied by Melford.

##### sfGFP gene sequence

ATGGTTAGCAAAGGTGAAGAACTGTTTACCGGCGTTGTGCCGATTCTGGTGGAAC TGGATGGTGATGTGAATGGCCATAAATTTAGCGTTCGTGGCGAAGGCGAAGGTG ATGCGACCAACGGTAAACTGACCCTGAAATTTATTTGCACCACCGGTAAACTGCC GGTTCCGTGGCCGACCCTGGTGACCACCCTGACCTATGGCGTTCAGTGCTTTAGC CGCTATCCGGATCATATGAAACGCCATGATTTCTTTAAAAGCGCGATGCCGGAAG GCTATGTGCAGGAACGTACCATTAGCTTCAAAGATGATGGCACCTATAAAACCC GTGCGGAAGTTAAATTTGAAGGCGATACCCTGGTGAACCGCATTGAACTGAAAG GTATTGATTTTAAAGAAGATGGCAACATTCTGGGTCATAAACTGGAATATAATTT CAACAGCCATAATGTGTATATTACCGCCGATAAACAGAAAAATGGCATCAAAGC GAACTTTAAAATCCGTCACAACGTGGAAGATGGTAGCGTGCAGCTGGCGGATCA TTATCAGCAGAATACCCCGATTGGTGATGGCCCGGTGCTGCTGCCGGATAATCAT TATCTGAGCACCCAGAGCGTTCTGAGCAAAGATCCGAATGAAAAACGTGATCAT ATGGTGCTGCTGGAATTTGTTACCGCCGCGGGCATTACCCACGGTATGGATGAAC TGTATAAAGGCAGCCACCATCATCATCACCATTAA

LifeAct-sfGFP gene sequence

TCCGCCCCATTGACGCAAATGGGCGGTAGGCGTGTACGGTGGGAGGTCTATATA AGCAGAGCTGGTTTAGTGAACCGTCAGATCCGCTAGCGCCACCATGGGCGTGGC CGACTTGATCAAGAAGTTCGAGTCCATCTCCAAGGAGGAGGGGGATCCACCGGT CGCCACCATGGTGAGCAAGGGCGAGGAGCTGTTCACCGGGGTGGTGCCCATCCT GGTCGAGCTGGACGGCGACGTAAACGGCCACAAGTTCAGCGTGCGCGGCGAGGG CGAGGGCGATGCCACCAACGGCAAGCTGACCCTGAAGTTCATCTGCACCACCGG CAAGCTGCCCGTGCCCTGGCCCACCCTCGTGACCACCCTGACCTACGGCGTGCAG TGCTTCAGCCGCTACCCCGACCACATGAAGCGCCACGACTTCTTCAAGTCCGCCA TGCCCGAAGGCTACGTCCAGGAGCGCACCATCAGCTTCAAGGACGACGGCACCT ACAAGACCCGCGCCGAGGTGAAGTTCGAGGGCGACACCCTGGTGAACCGCATCG AGCTGAAGGGCATCGACTTCAAGGAGGACGGCAACATCCTGGGGCACAAGCTGG AGTACAACTTCAACAGCCACAACGTCTATATCACCGCCGACAAGCAGAAGAACG GCATCAAGGCCAACTTCAAGATCCGCCACAACGTGGAGGACGGCAGCGTGCAGC TCGCCGACCACTACCAGCAGAACACCCCCATCGGCGACGGCCCCGTGCTGCTGC CCGACAACCACTACCTGAGCACCCAGTCCGTGCTGAGCAAAGACCCCAACGAGA AGCGCGATCACATGGTCCTGCTGGAGTTCGTGACCGCCGCCGGGATCACTCACG GCATGGACGAGCTGTACAAGTAAGCGGCCGCGACTCTAGATCATAATCAGCCATACCACATTTGTAGAGGTTTTACTTGCTTTAAAAAACCTCCCACACCTCCCCCTGA ACCTGAAACATAAAATGAA

## Supporting Tables

**Table S1.**
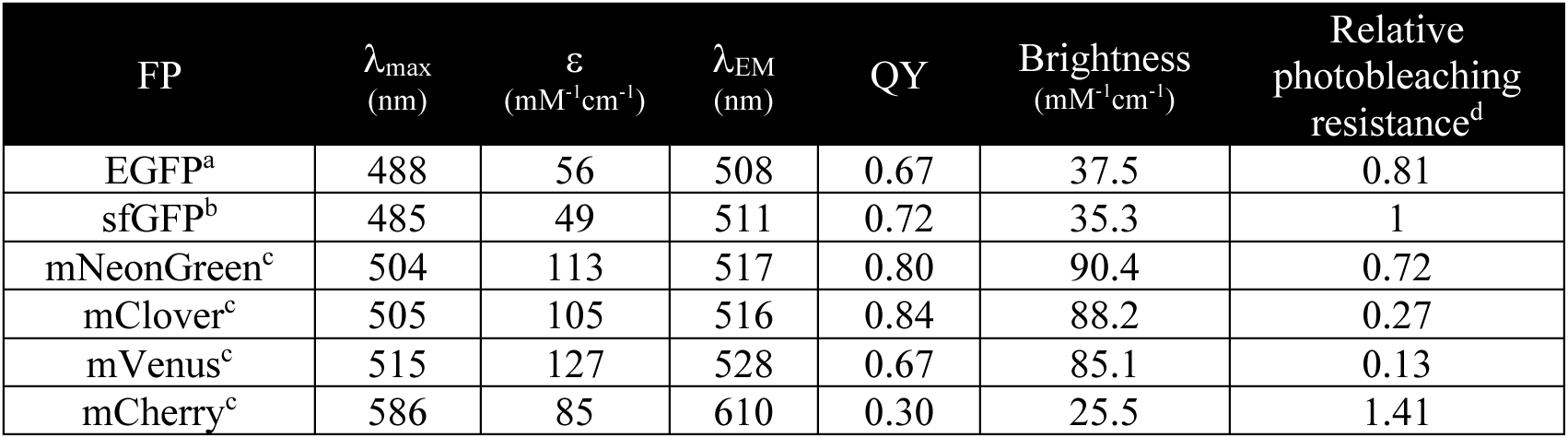
Comparison of different fluorescent proteins. ^a^, from FPBase (https://www.fpbase.org) ^1^. ^b^, from Reddington et al ^2^. ^c^, from the Cranfill et al ^3^ ^d^, based on t (1/2) values published by Cranfill et al ^3^. Values relative to sfGFP, which is set to 1.

**Table S2.**
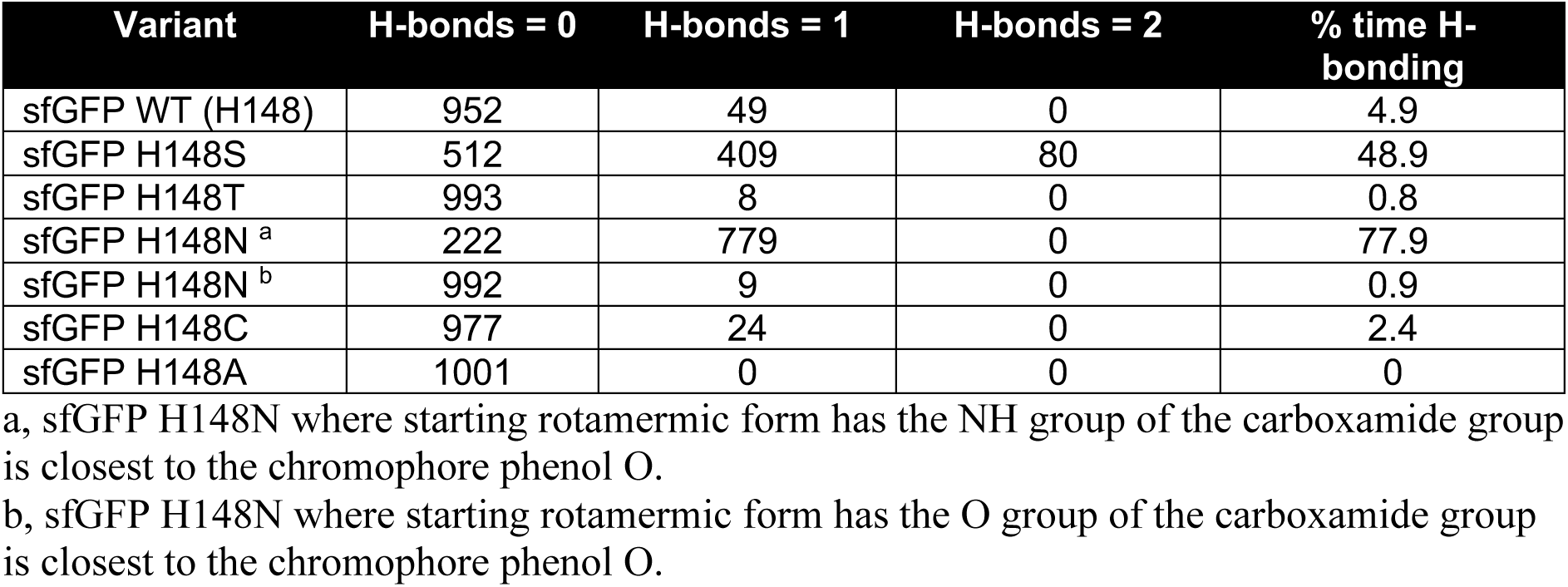
H-bond frequency between chromophore and residue 148 over 10ns simulations.

**Table S3.**
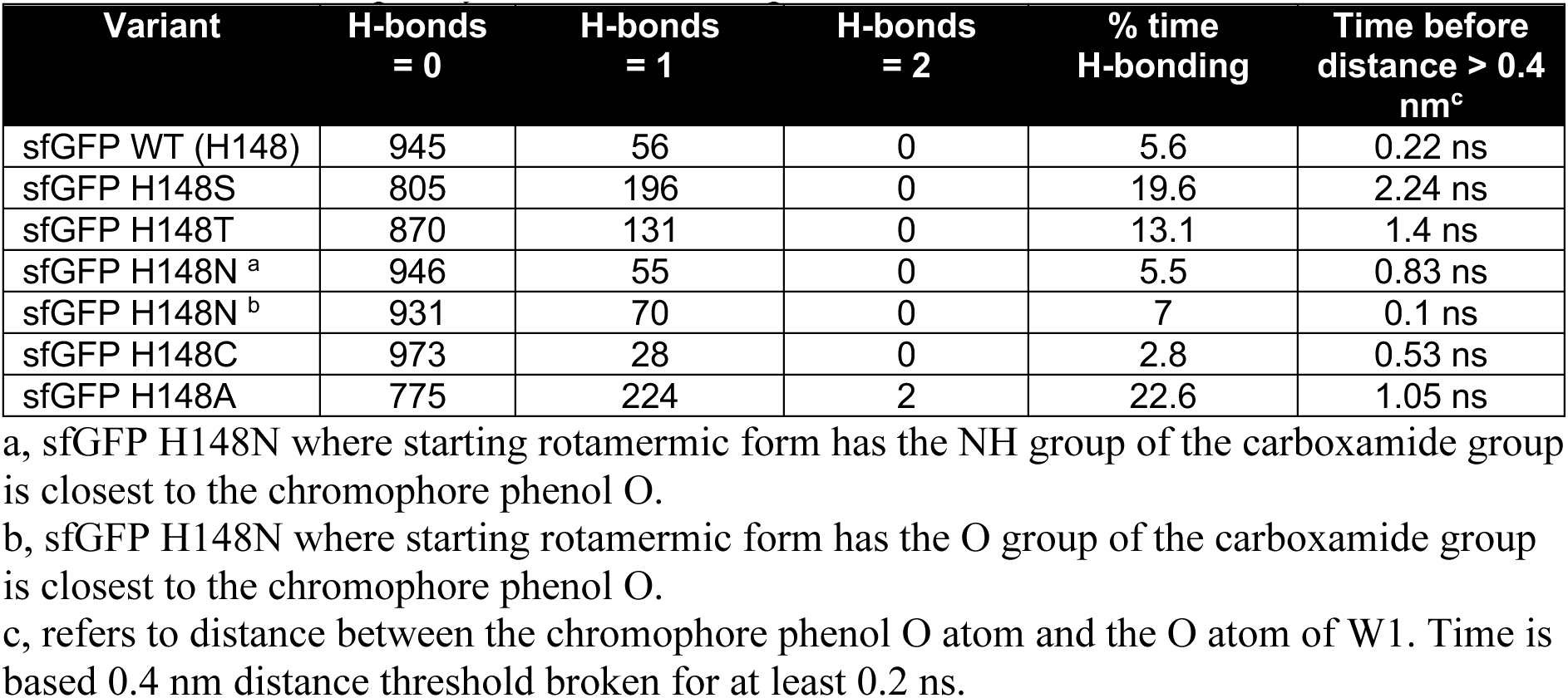
H-bond frequency between chromophore and W1 water over 10ns simulations.

**Table S4.**
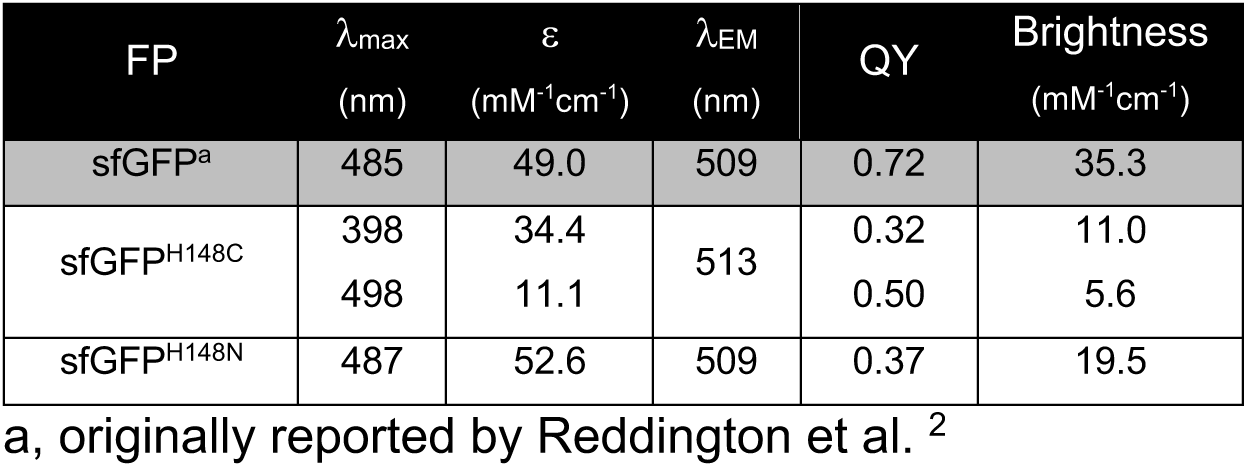
Spectral properties of sfGFP-H148X variants.

**Table S5.**
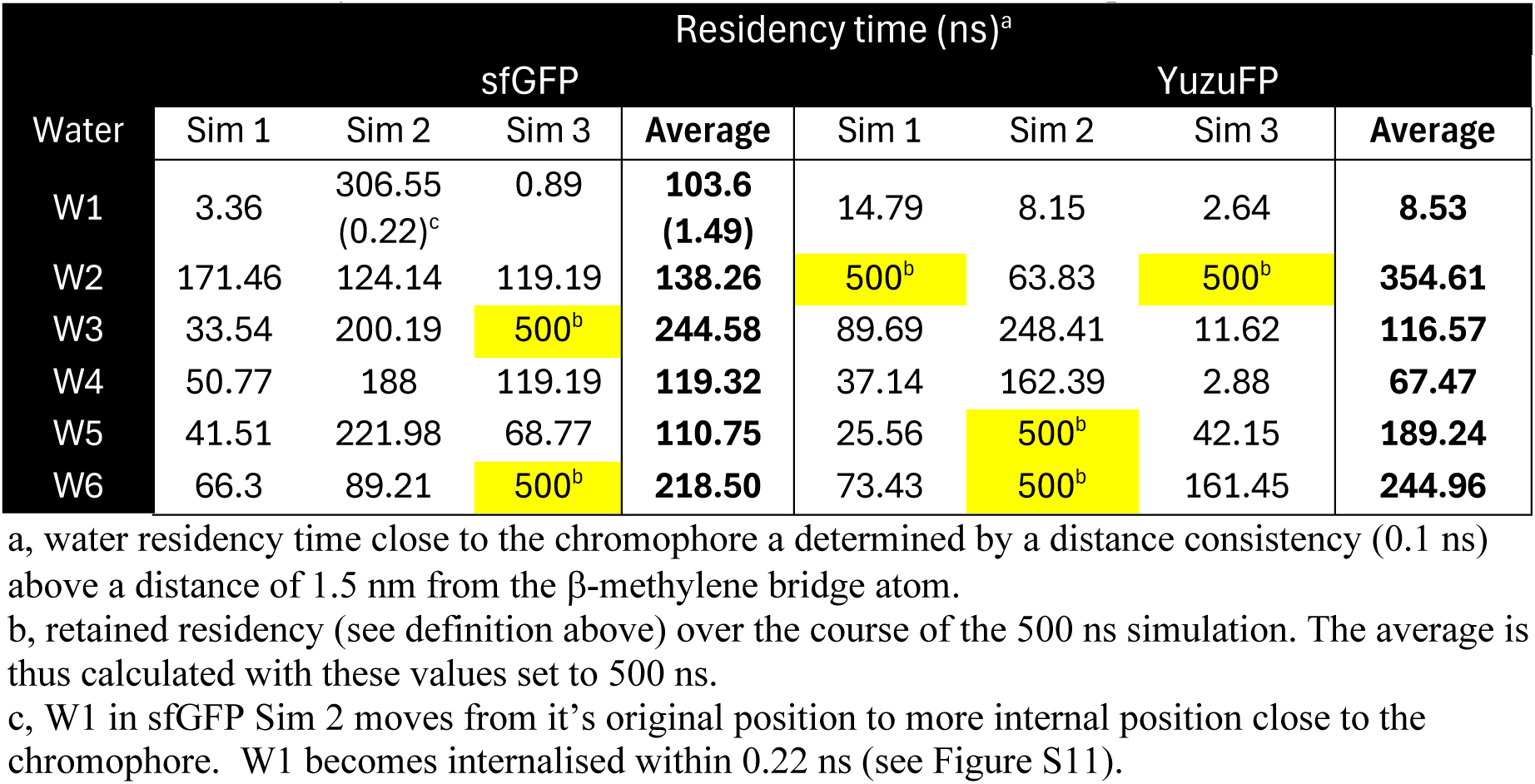
Residency time of water molecules close to the chromophore.

**Table S6.**
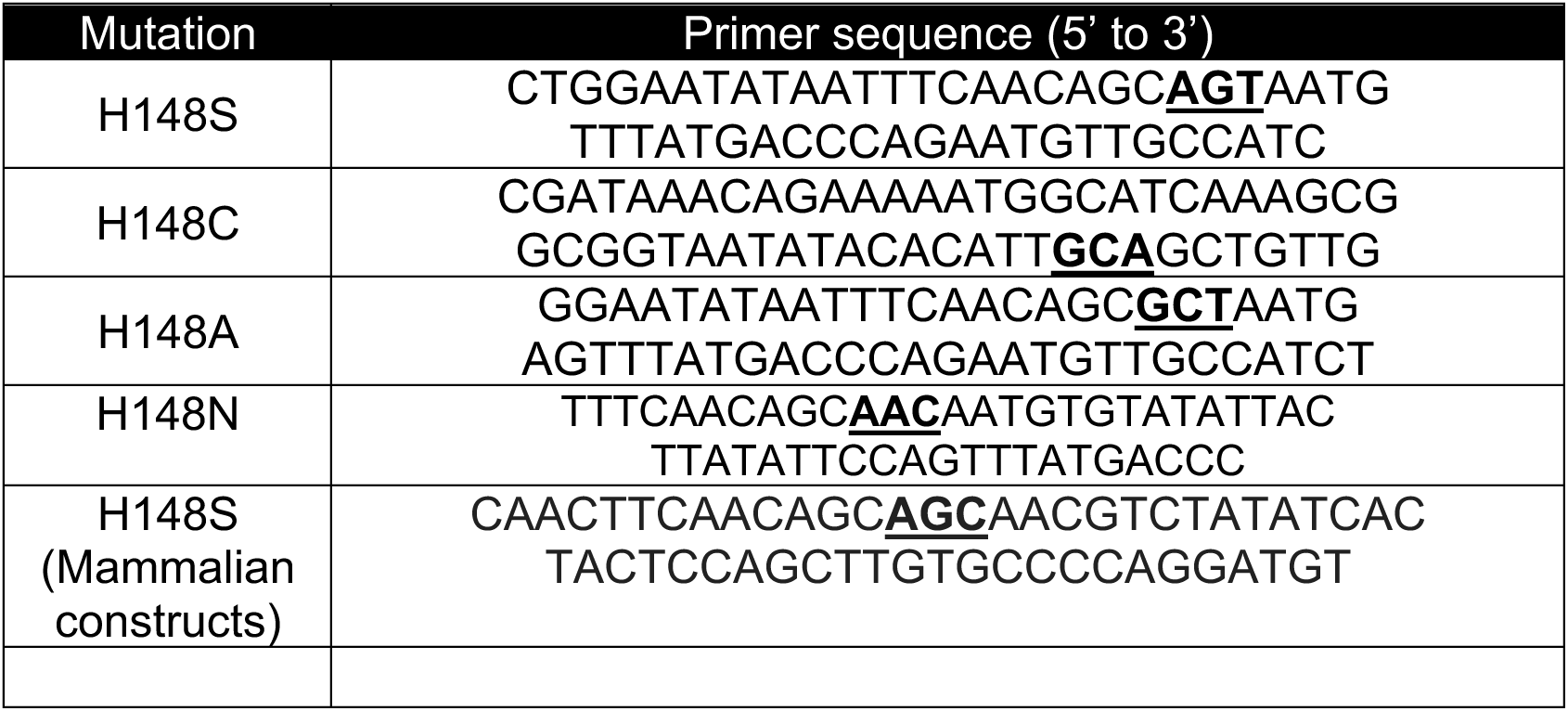
Primer combinations and sequences for introducing H148X mutations. Bases in bold and underlined represent the mutation. Top primer is the “forward” and bottom the “reverse” primer in a whole plasmid PCR.

## Supporting Figures

**Figure S1.**
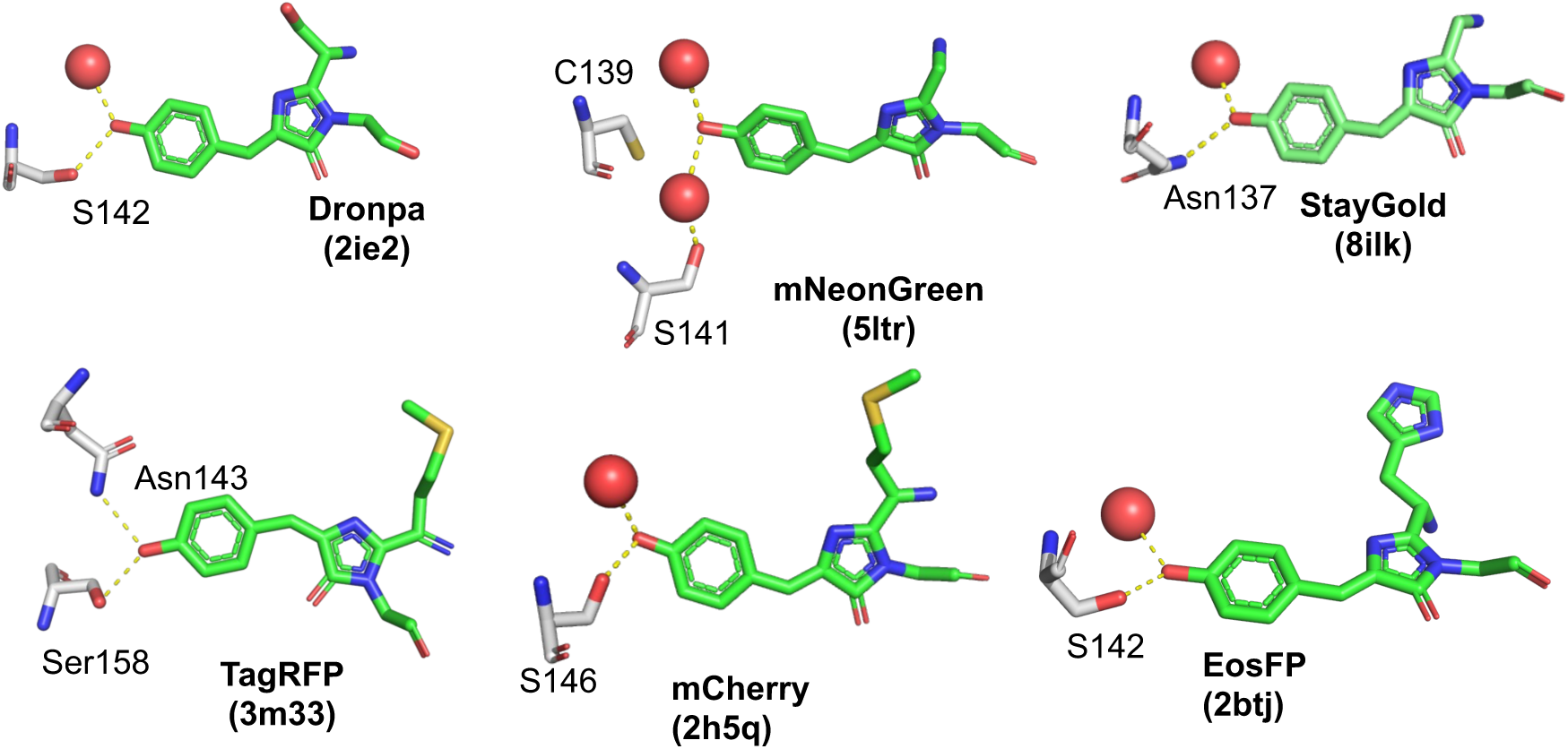
Additional examples of residues equivalent to H148 that interact with the chromophore phenol group: Dronpa ^4^ (PDB 2ie2), mNeonGreen ^5^(PDB 5ltr), StayGold^6^ (PDB 8ilk), TagRFP ^7,8^ (PDB 3m33), mCherry ^9^(PDB 2h5q), EosFP ^10^ (PDB 1zux). In all cases the chomophore is coloured green, interacting residues equivalent to H148 in sfGFP grey and water molecules are red spheres.

**Figure S2.**
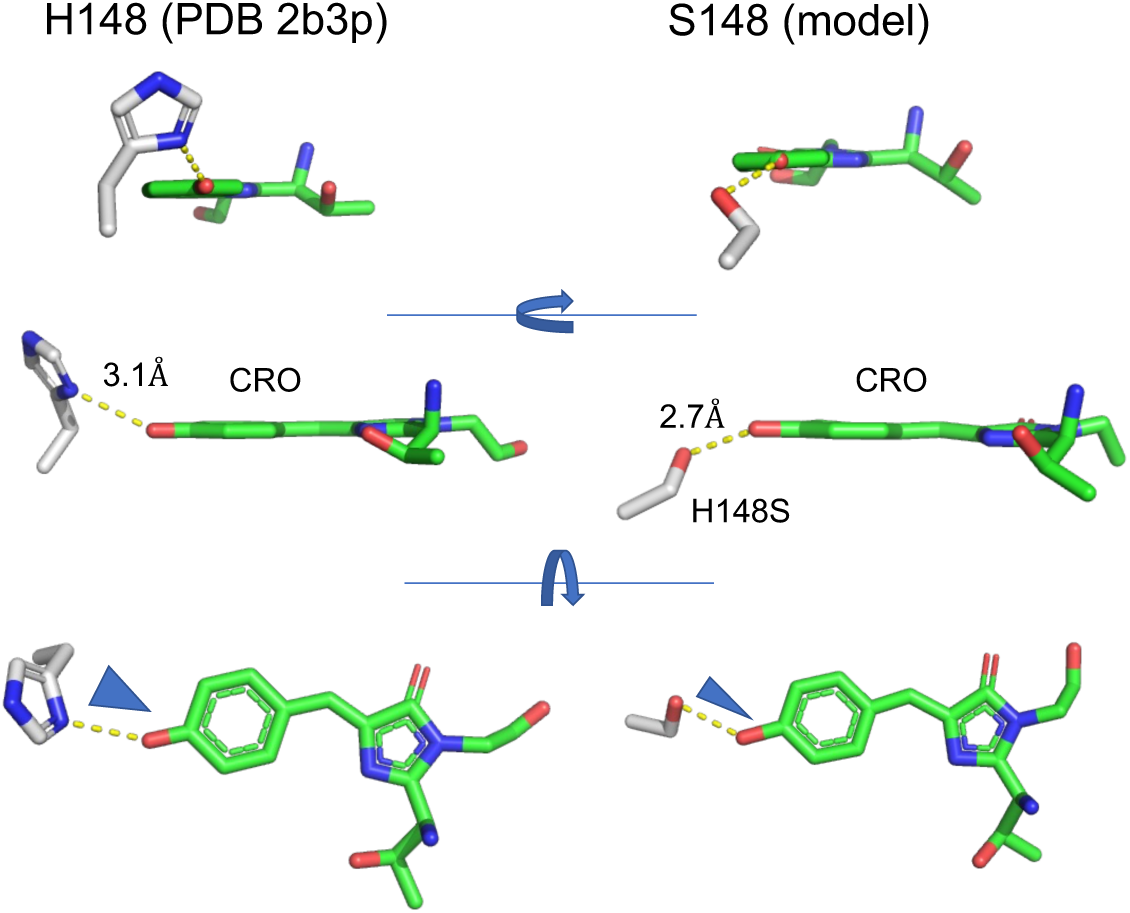
Relative conformations of residue 148 (grey sticks) and chromophore (green sticks). Left hand side is the position of H148 taken from the known structure of sfGFP (2b3p.pdb ^11^). Modelled structure of the H148S is the clustered average of individual MD trajectory outputs after 10 ns of molecular dynamics.

**Figure S3.**
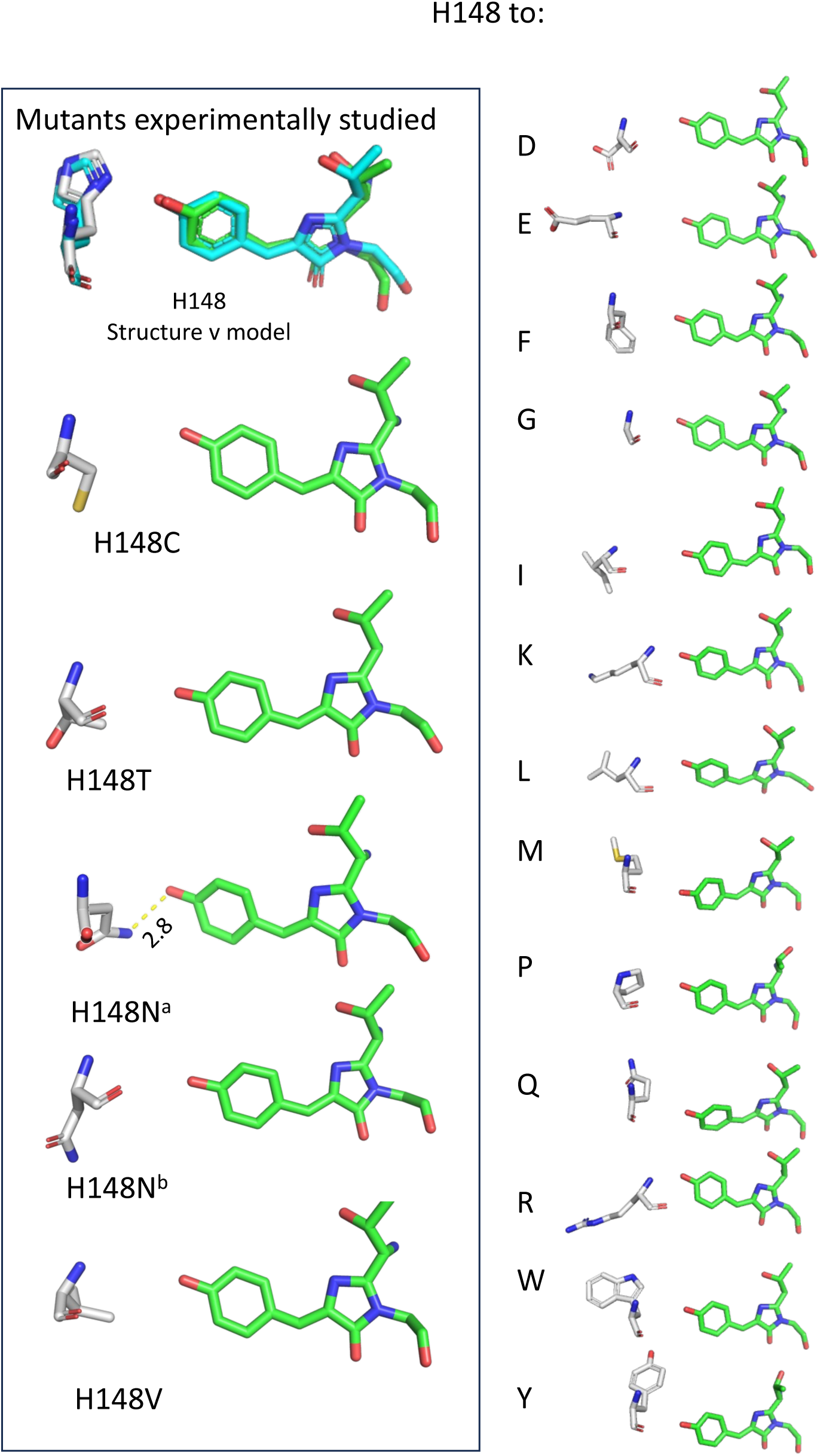
Modelling the H148X mutations. Modelled structures of the H148X mutations are the clustered average of individual MD trajectory outputs after 10 ns of molecular dynamics. Mutations on the left hand side have been experimentally analysed. For H148N, two model outcomes are shown: H148N^a^ is from a starting point of the NH group of the carboxamide group closest to the chromophore phenol O; H148N^b^ is from a starting of the O group of the carboxamide group closest to the chromophore phenol O.

**Figure S4.**
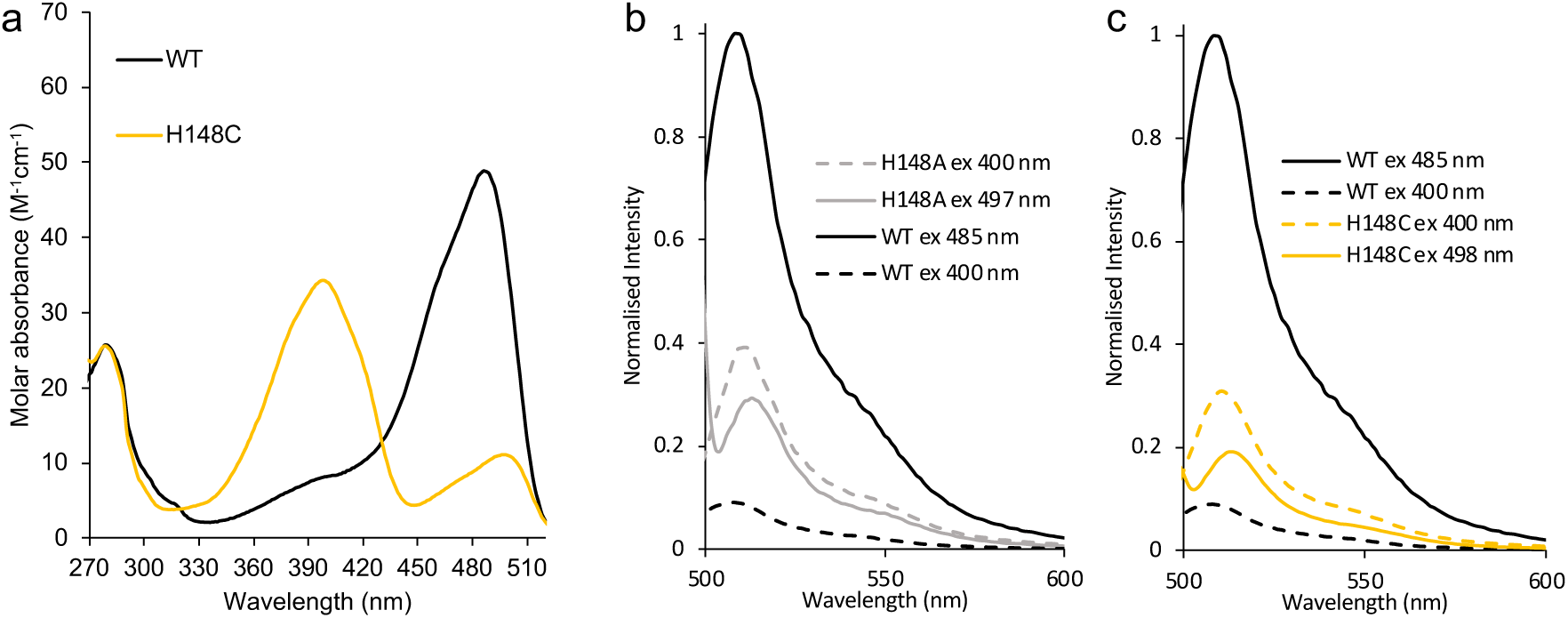
Absorbance and Fluorescence emission spectra for sfGFP variants. (a) Absorbance spectra for sfGFP WT (black) and H148C (orange). Fluorescence emission spectra for (b) H148A and (c) H148C. For clarity, the same WT sfGFP spectra are plot in each of the 4 panels and coloured black. Excitation at 485-498 nm is shown as solid lines and excitation at 400 nm as dashed lined. (b) H148A is coloured grey with excitation at 497 nm shown as solid lines and excitation at 400 nm as dashed lined. (c) H148C is coloured orange with excitation at 498 nm shown as solid lines and excitation at 400 nm as dashed lined.

**Figure S5.**
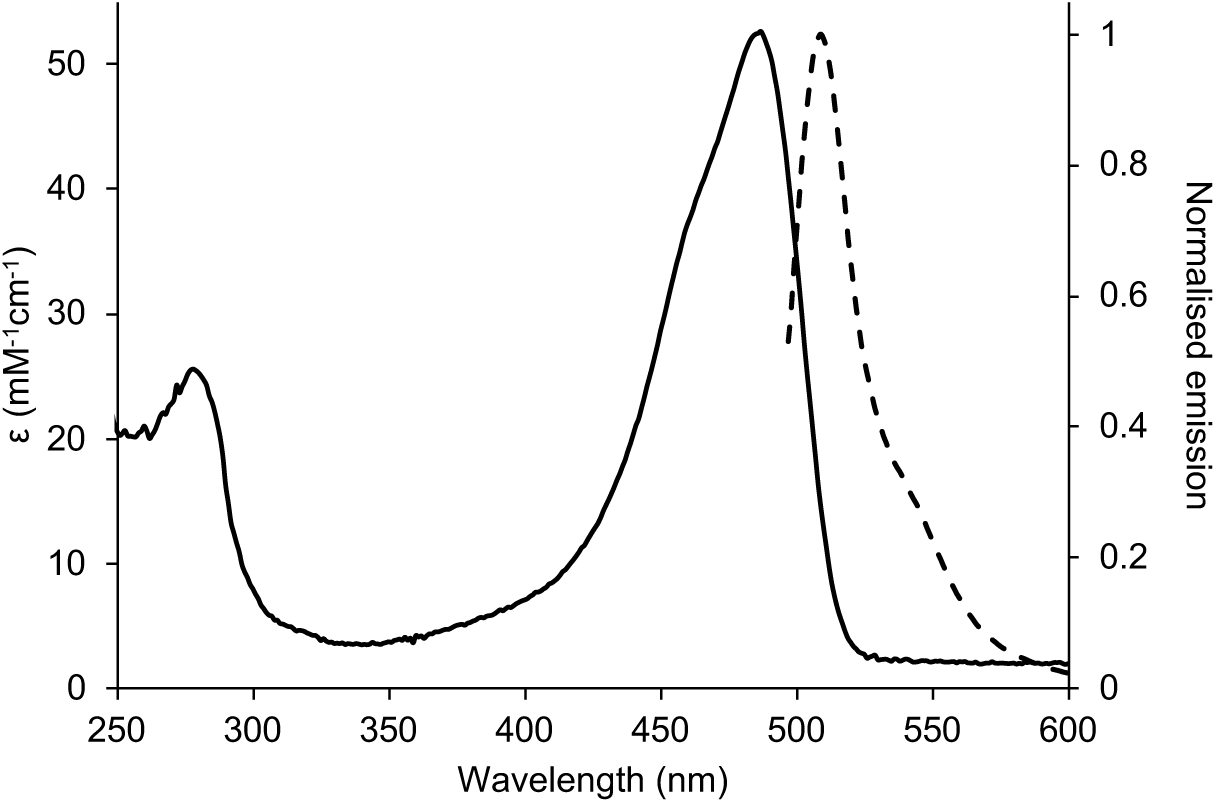
Absorbance and fluorescence spectra of sfGFP-H148N. The absorbance spectrum are shown as a solid line and normalised to the molar absorbance coefficient. Fluorescence emission spectra on excitation at 490 nm is shown as dashed lined and normalised to 1 based on the maximum emission intensity at 509 nm.

**Figure S6.**
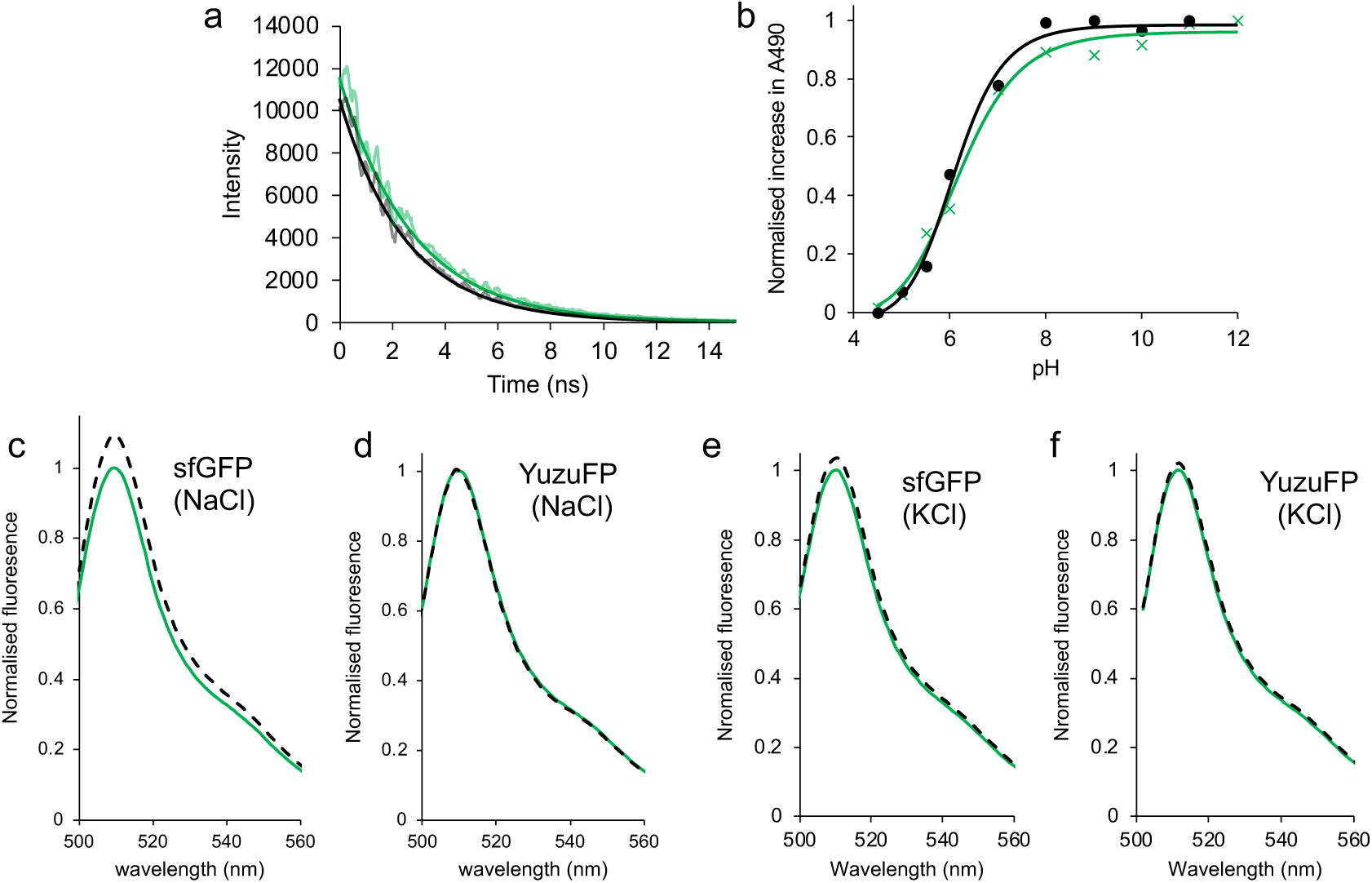
Analysis of YuzuFP. (a) The fluorescence intensity decay curve of YuzuFP (green) and sfGFP (black) on excitation at 473 nm. Data where fit to a single exponential decay in Graphpad Prism. (b) pKa fo YuzuFP (green) and sfGFP (black) as measured by the increase in absorbance at 490 nm. Data where fit to a least squares fit sigmodal model in Graphpad Prism. (c-f) The effect of 150 mM salt (either NaCl or KCl) on fluorescence emission, as indicated in the figure. Green solid lines represent protein in 50 mM Tris pH 8.0 and dashed black lines in the presence of 150 mM salt.

**Figure S7.**
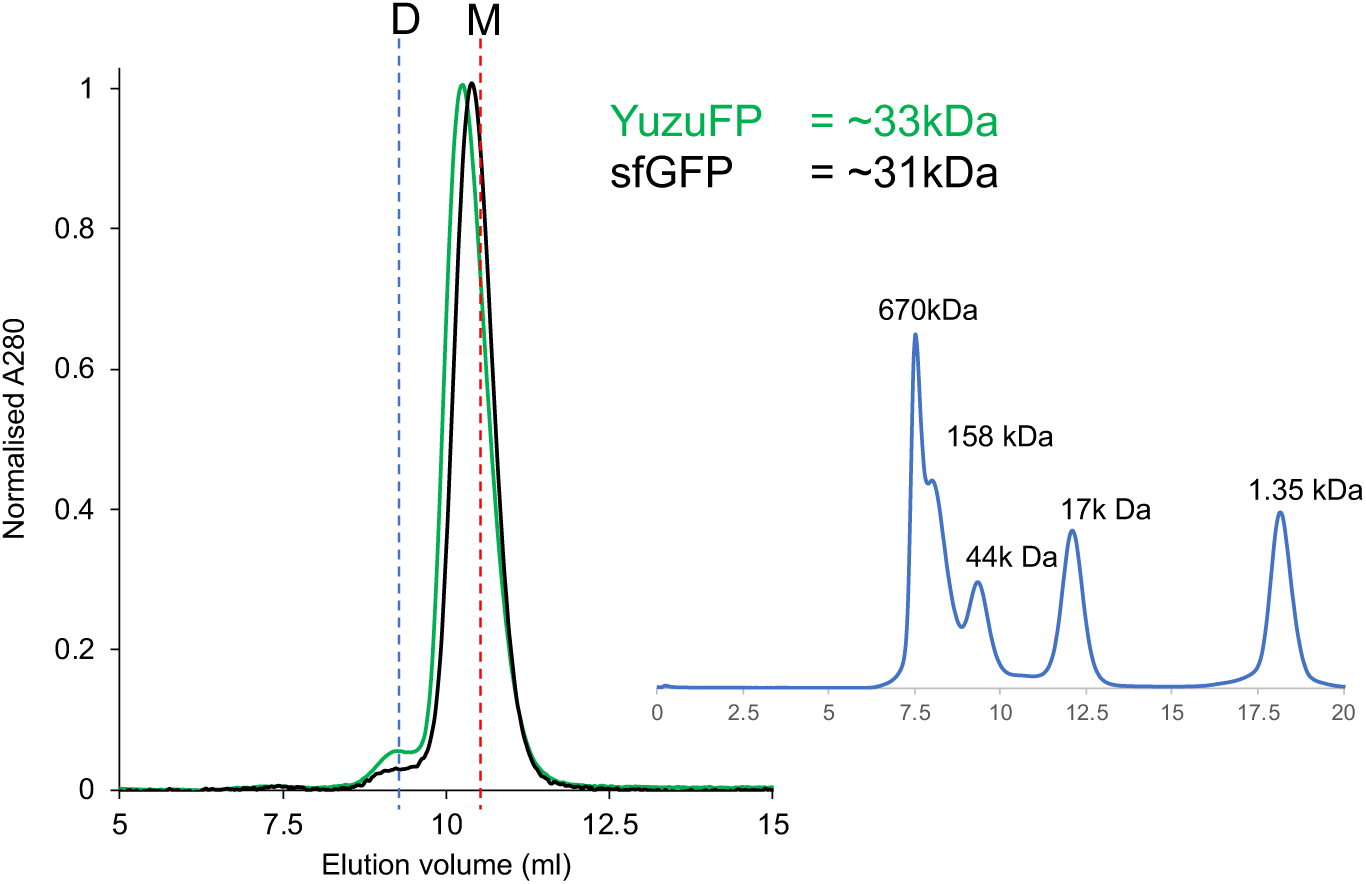
Analysis of quaternary structure YuzuFP by size exclusion chromatography (SEC). The black and green lines represent 50 μM sfGFP and 50 μM YuzuFP, respectively. SEC was performed using a calibrated Superdex 75 10/300GL column (inset). Estimated elution volumes for a theoretical monomer (M; 27 kDa) and dimer (D; 54 kDa) are shown by red and blue sashed lines, respectively. YuzuFP eluted very slightly earlier than sfGFP, giving an estimated molecular weight of ∼33 kDa compared to 31 kDa for sfGFP. A small shoulder at ∼9.3 ml suggests a small amount of dimer (∼5%) may be present for both sfGFP and YuzuFP.

**Figure S8.**
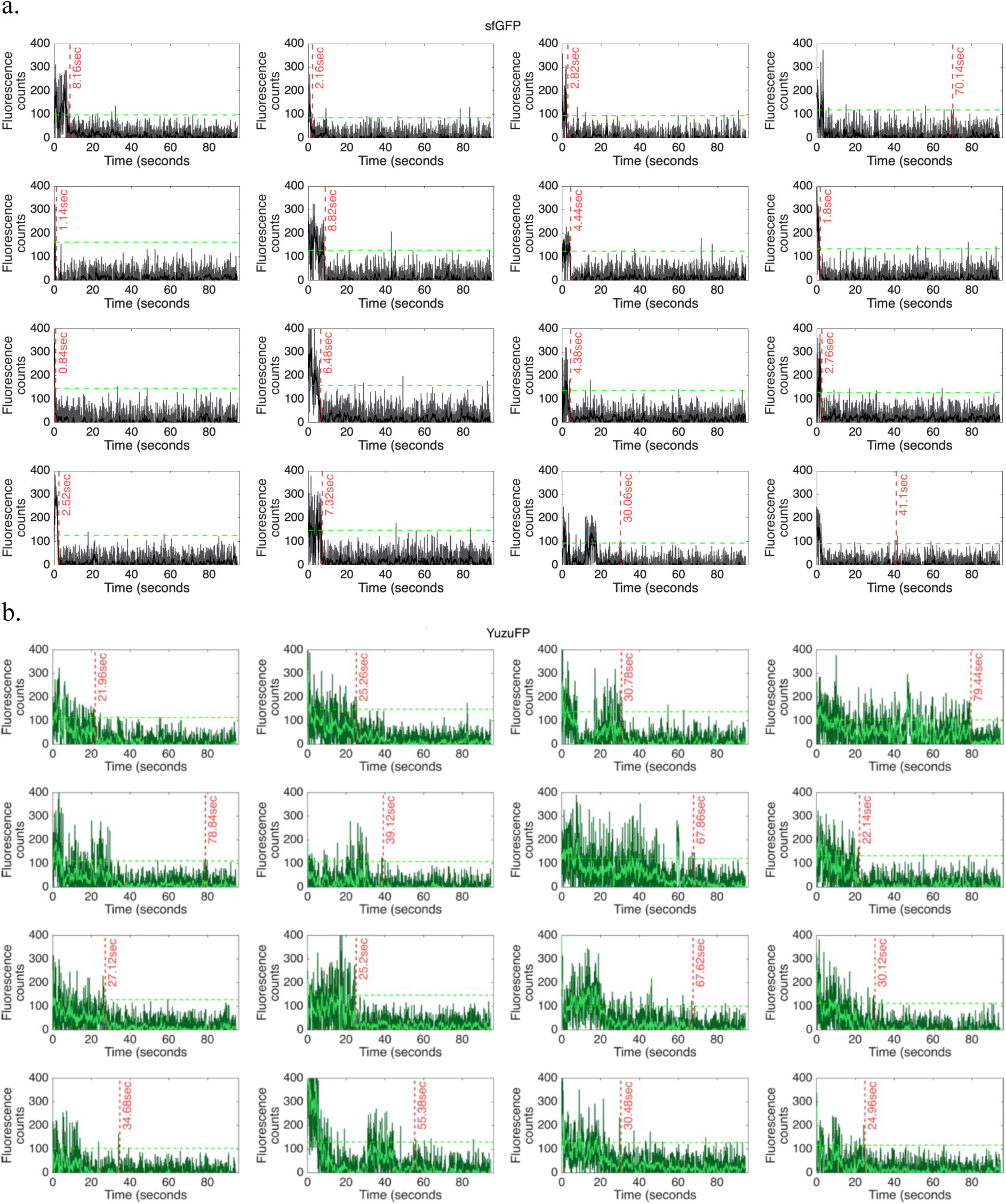
Exemplar single molecules fluorescence traces for (a) sfGFP and (b) YuzuFP. Single molecule data is extracted from a micrograph timeseries acquired using a total internal reflection fluorescence (TIRF) microscopy imaging system. Trace’s are generated from the mean intensities of 4 by 4-pixel regions of interest corresponding to individual fluorophores. Raw data (grey and dark green) are plotted along side data passed through a forward-backward moving window average filter (black and light green). In order to clearly identify “on” and “off” fluorescence states thresholding of the raw data to 10 standard deviations beyond the mean background was used (green dashed line). Using this in combination with a temporal threshold allowed for identification of photobleaching lifetimes of all single molecules (red dashed lines and times in seconds).

**Figure S9.**
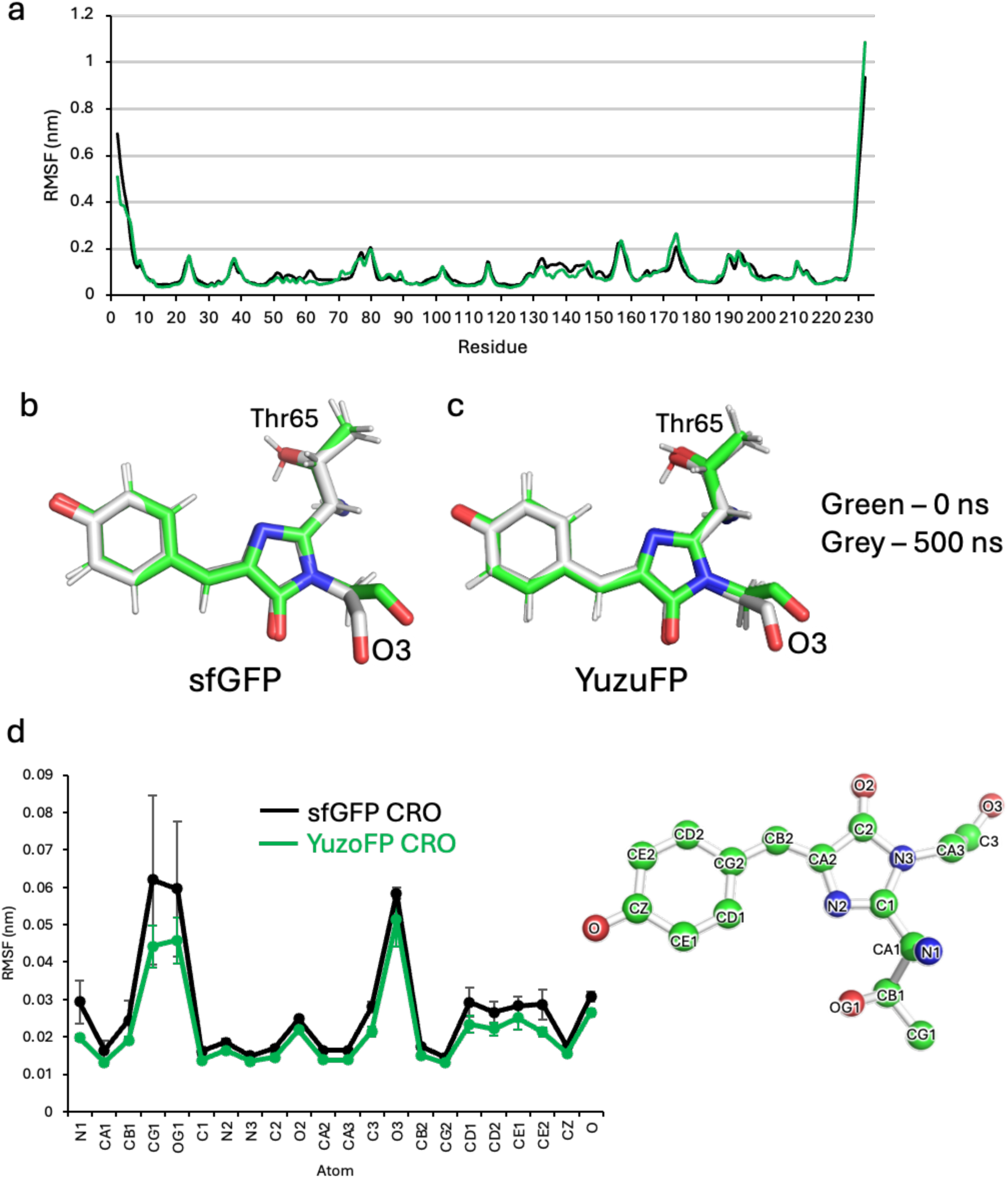
Change in backbone and chromophore structure over the course of MD simulation. (a) Per residue backbone (Cα) RMSF of sfGFP (black) and YuzuFP (green). RMSF values shown are an average of the 3 independent simulations. The chromophore structure at the start (green, 0 ns) and end (grey; 500 ns) of a single simulation for (b) sfGFP and (c) YuzuFP. (d) The root mean square fluctuation (RMSF) of each of the chromophore heavy atoms (as annotated in the molecular structure to the right) for sfGFP (black) and YuzuFP (green). The error bars represent the standard deviation between the 3 independent 500 ns simula

**Figure S10.**
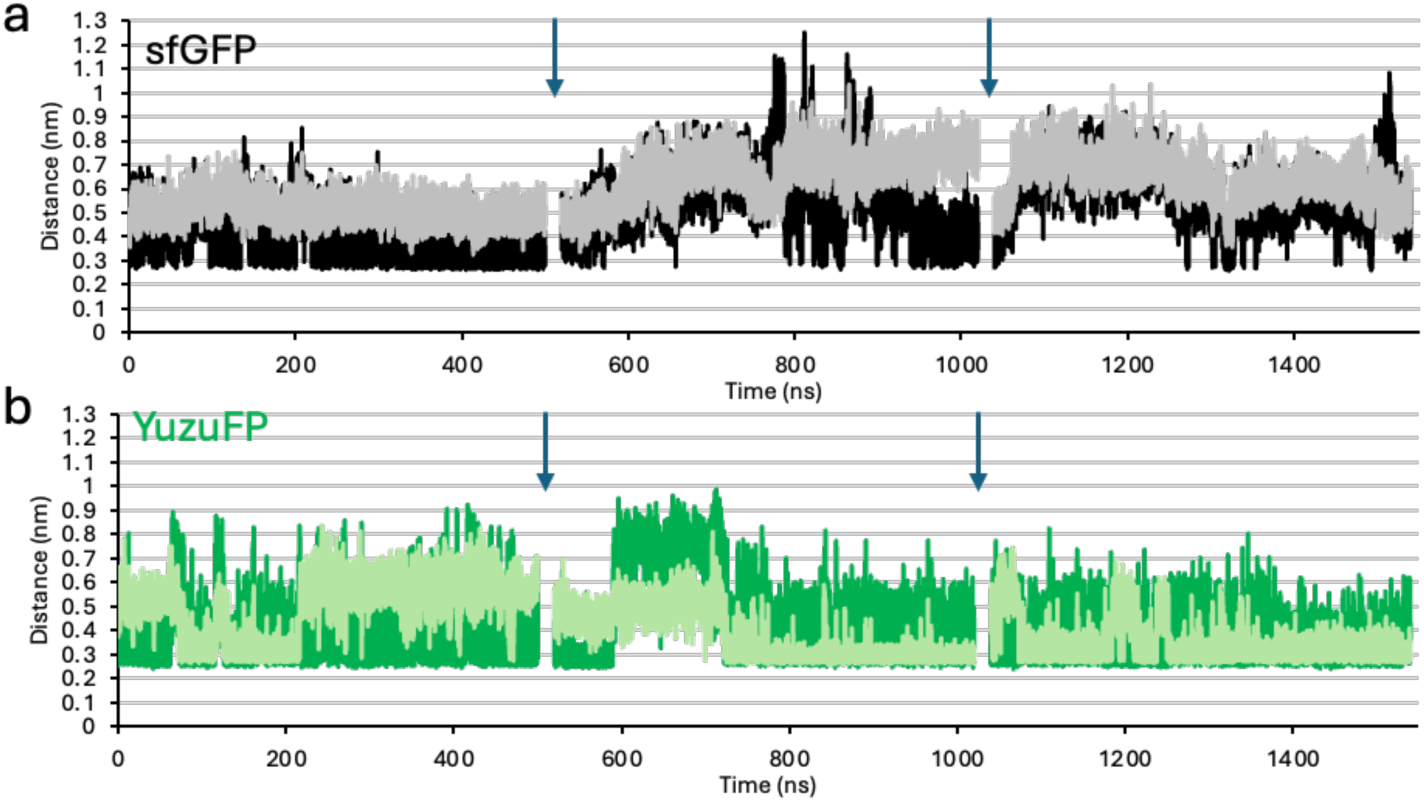
The pair-wise distance in (a) sfGFP and (b) YuzuFP, between chromophore phenol oxygen and residue 148 backbone and side chain H-bond donor heavy atoms. In (a), the backbone amide nitrogen is coloured grey and the side chain imidazole nitrogen of H148 is coloured black. In (b), the backbone amide nitrogen is coloured lime and the side chain hydroxyl oxygen of S148 is coloured green. The plots are a concatenation of the three 500 ns MD simulations, where the down arrows show the separation between each simulation.

**Figure S11.**
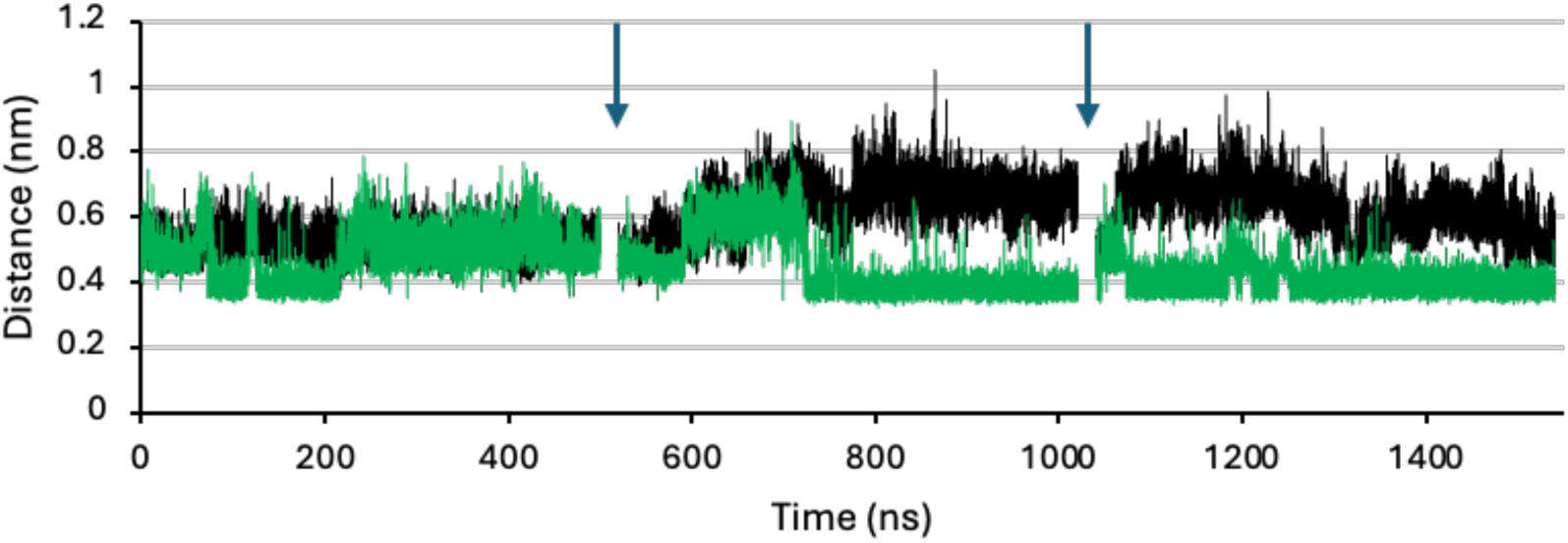
The distance between the chromophore phenol oxygen and the Cα of residue 148 in sfGFP (black) and YuzuFP (green). Each plot is a concatenation of 3 separate 500 ns simulations. The start and end of each simulation is shown by the down arrows.

**Figure S12.**
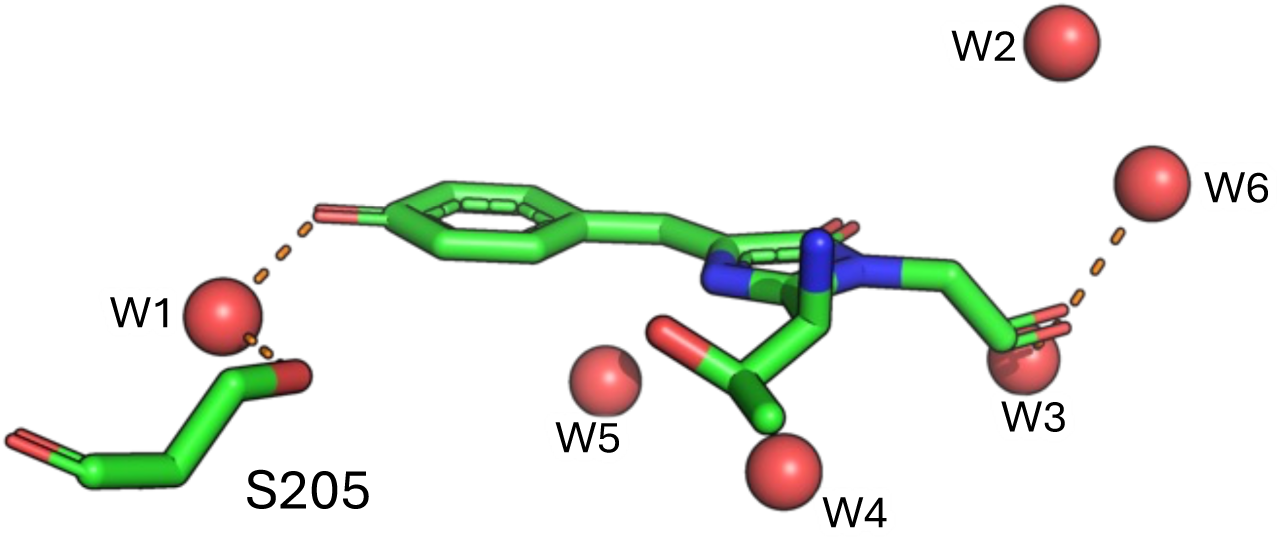
Water molecules within proximity of the chromophore. Red spheres are water molecules with the W1 water molecule labelled. Dashed lines are polar interactions between the chromophore and water molecules. The structure shown is PDB 2b3p.

**Figure S13.**
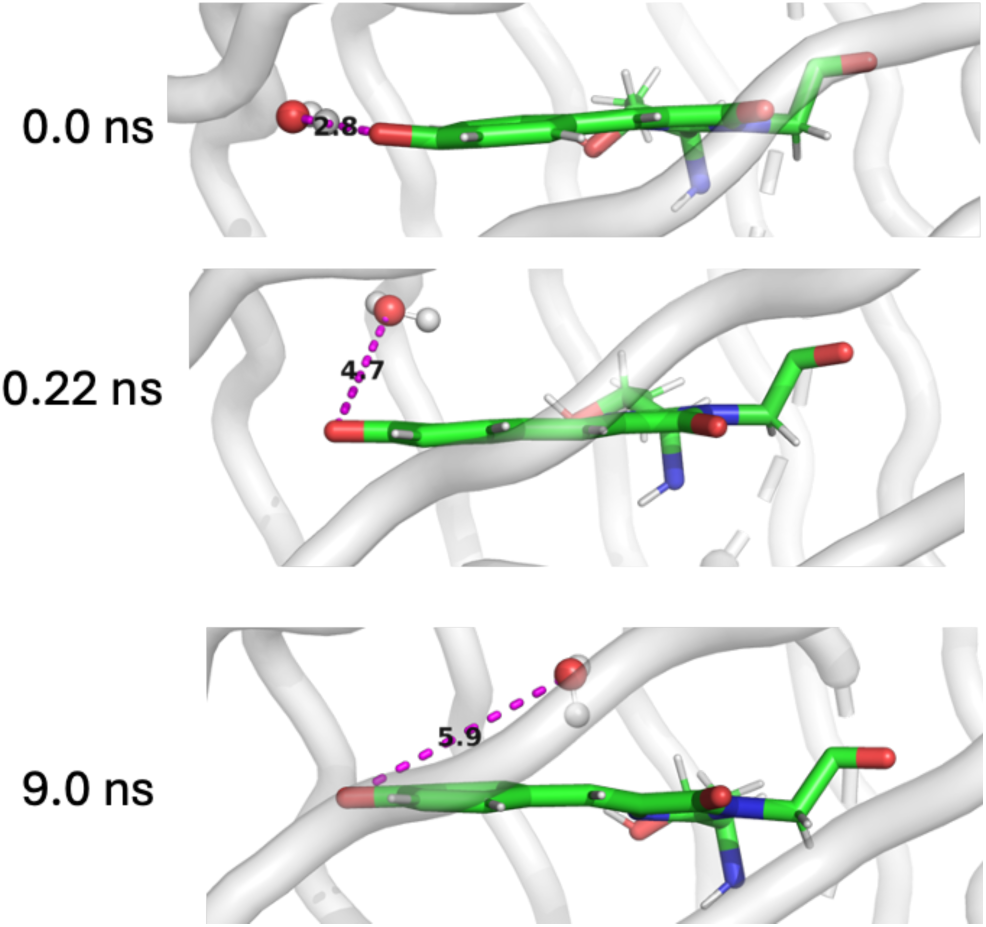
Position of water molecule W1 in sfGFP simulation 2. Within 0.22ns, W1 is internalised and no longer able to form a H-bond with the chromophore phenol oxygen. The chromophore is coloured green and water molecule W1 shown as balled and stick. The dashed magenta lines and associated numeric value represent the distance between the chromophore oxygen and the W1 water molecule (in Ångstroms).

**Figure S14.**
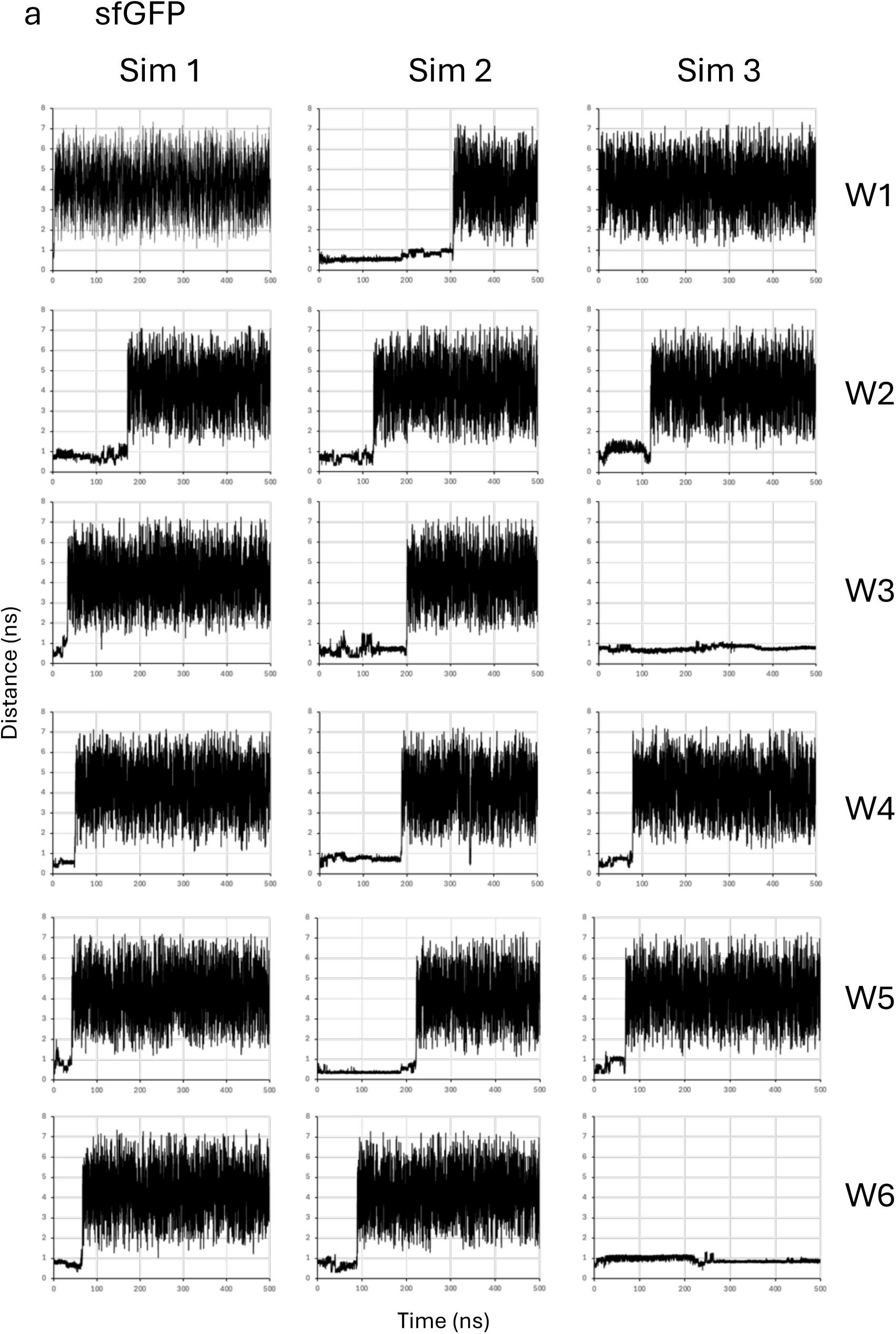
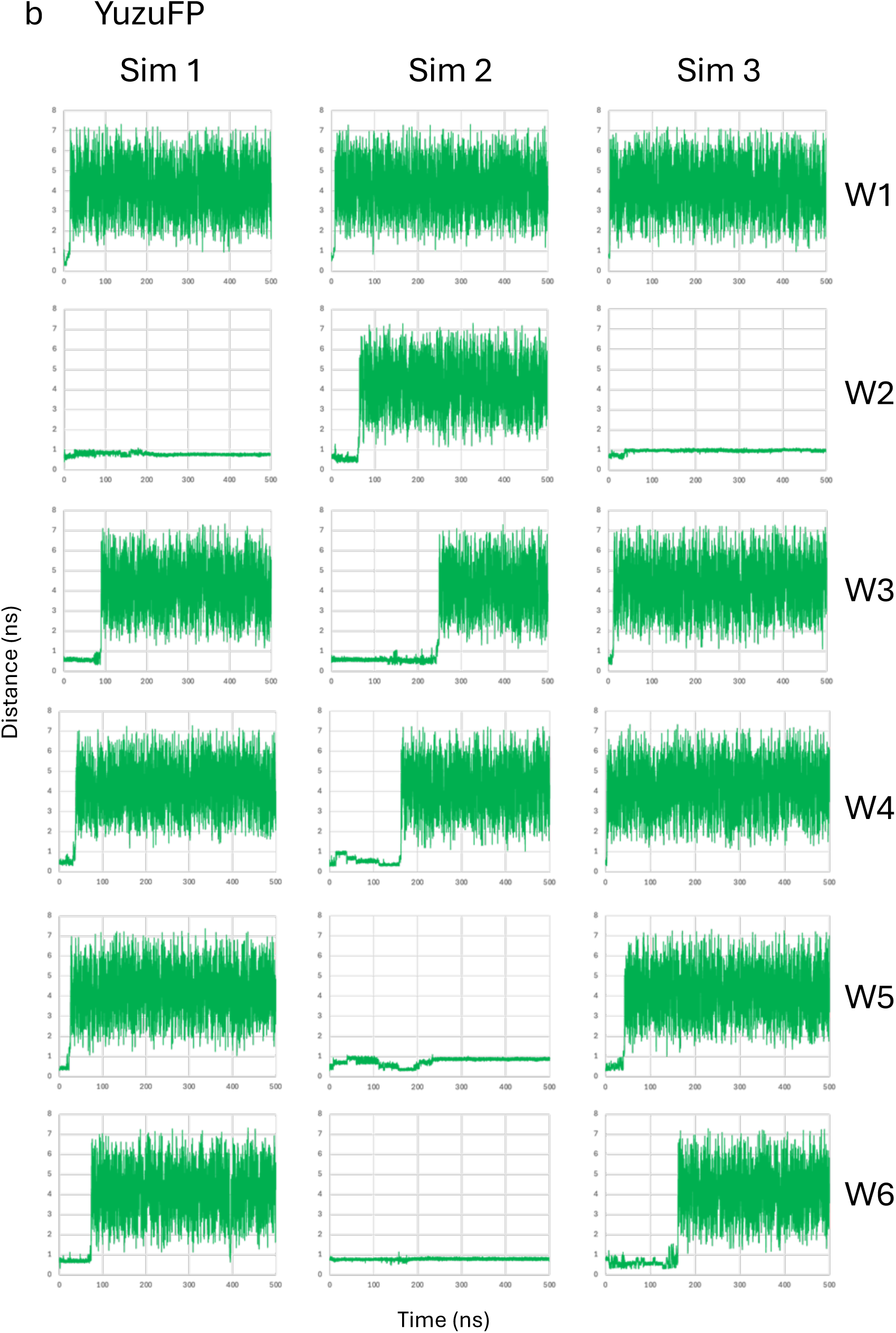
The pairwise distance between chromophore (atom 967 as part of the β-methylene bridge) and individual water oxygen atom (W1 to W6) from each 500 ns simulation (Sim 1 to 3) in (a) sfGFP and (b) YuzoFP. In sfGFP, W1 to W6 corresponding to oxygen atom IDs 3642, 3618, 3624, 3663, 3708, 3609, respectively. For YuzoFP W1 to W6 corresponding to oxygen atom IDs 3636, 3612, 3618, 3657, 3702, 3603.

**Figure S15.**
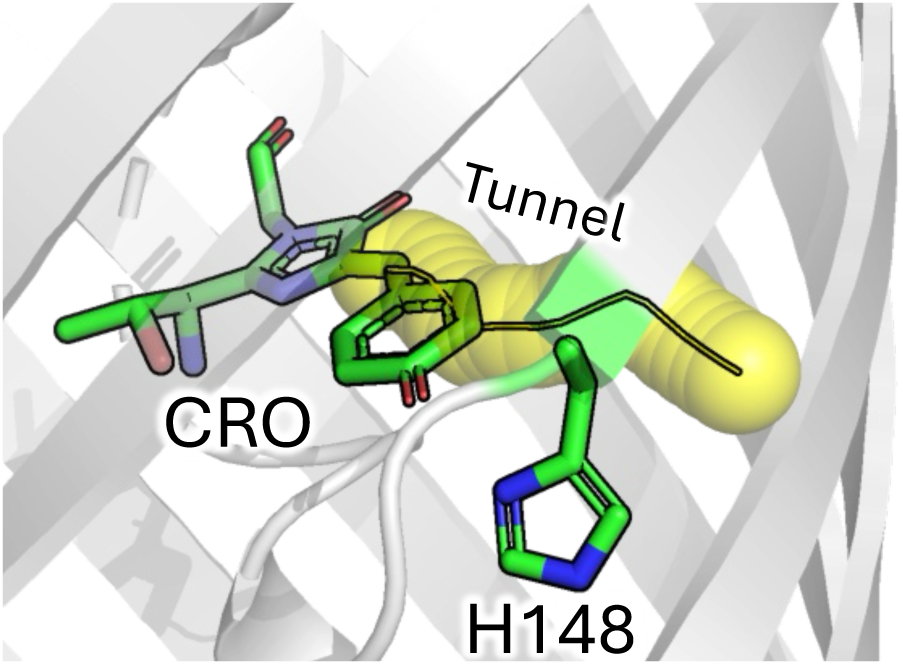
Tunnel (yellow; 1.1 Å probe radius) between CRO and solvent as calculated using CAVER ^12,13^ and the original PDB for sfGFP (2b3p)^11^.

