## Supplementary methods, Tables and figures for "Molecular dynamics guided engineering of *Aequorea victoria* Green Fluorescent Protein chromophore interactions generates a brighter variant with improved photobleaching resistance"

**Supplementary Information.**

**Supporting Methods.**

**Autoinduction media**

The following lists the auto-induction media used to produce sfGFP proteins present in the pBAD plasmid: 1% (w/v) tryptone; 0.5 %(w/v) yeast extract; 0.5% (v/v) glycerol; 0.05 % (w/v) glucose; 0.2% lactose; 25 mM $Na_2HPO_4$; 25 mM $KH_2PO_4$; 50 mM $NH_4Cl$; 5 mM $NaSO_4$; 2 mM $MgSO_4$; 1 x trace metals (4 µM $CaCl_2$; 2 µM, $MnCl_2$, 2 µM $ZnSO_4$, 0.4 µM $CoCl_2$, 0.4 µM $CuCl_2$, 0.4 $NiCl_2$, 0.4 uM $Na_2MoO_4$, 0.4 µM $H_3BO_3$ and 10 µM $FeCl_3$ in ultra-pure water) and 0.05 % (w/v) L-arabinose. The medium was supplemented with 50 µg/ml ampicillin. All chemicals were supplied by Melford.

**sfGFP gene sequence.**

ATGGTTAGCAAAGGTGAAGAACTGTTTACCGGCGTTGTGCCGATTCTGGTGGAACTGGATGGTGAT
GTGAATGGCCATAAATTTAGCGTTCGTGGCGAAGGCGAAGGTGATGCGACCAACGGTAAACTGACC
CTGAAATTTATTTGCACCACCGGTAAACTGCCGGTTCCGTGGCCGACCCTGGTGACCACCCTGACCT
ATGGCGTTCAGTGCTTTAGCCGCTATCCGGATCATATGAAACGCCATGATTTCTTTAAAAGCGCGAT
GCCGGAAGGCTATGTGCAGGAACGTACCATTAGCTTCAAAGATGATGGCACCTATAAAACCCGTGC
GGAAGTTAAATTTGAAGGCGATACCCTGGTGAACCGCATTGAACTGAAAGGTATTGATTTTAAAGA
AGATGGCAACATTCTGGGTCATAAACTGGAATATAATTTCAACAGCCATAATGTGTATATTACCGCC
GATAAACAGAAAAATGGCATCAAAGCGAACTTTAAAATCCGTCACAACGTGGAAGATGGTAGCGTG
CAGCTGGCGGATCATTATCAGCAGAATACCCCGATTGGTGATGGCCCGGTGCTGCTGCCGGATAAT
CATTATCTGAGCACCCAGAGCGTTCTGAGCAAAGATCCGAATGAAAAACGTGATCATATGGTGCTGC
TGGAATTTGTTACCGCCGCGGGCATTACCCACGGTATGGATGAACTGTATAAAGGCAGCCACCATCA
TCATCACCATTAA

LifeAct-sfGFP gene sequence

TCCGCCCCATTGACGCAAATGGGCGGTAGGCGTGTACGGTGGGAGGTCTATATAAGCAGAGCTGGT
TTAGTGAACCGTCAGATCCGCTAGCGCCACCATGGGCGTGGCCGACTTGATCAAGAAGTTCGAGTC
CATCTCCAAGGAGGAGGGGGATCCACCGGTCGCCACCATGGTGAGCAAGGGCGAGGAGCTGTTCA
CCGGGGTGGTGCCCATCCTGGTCGAGCTGGACGGCGACGTAAACGGCCACAAGTTCAGCGTGCGC
GGCGAGGGCGAGGGCGATGCCACCAACGGCAAGCTGACCCTGAAGTTCATCTGCACCACCGGCAA
GCTGCCCGTGCCCTGGCCCACCCTCGTGACCACCCTGACCTACGGCGTGCAGTGCTTCAGCCGCTAC
CCCGACCACATGAAGCGCCACGACTTCTTCAAGTCCGCCATGCCCGAAGGCTACGTCCAGGAGCGCA
CCATCAGCTTCAAGGACGACGGCACCTACAAGACCCGCGCCGAGGTGAAGTTCGAGGGCGACACCC
TGGTGAACCGCATCGAGCTGAAGGGCATCGACTTCAAGGAGGACGGCAACATCCTGGGGCACAAG
CTGGAGTACAACTTCAACAGCCACAACGTCTATATCACCGCCGACAAGCAGAAGAACGGCATCAAG

GCCAACTTCAAGATCCGCCACAACGTGGAGGACGGCAGCGTGCAGCTCGCCGACCACTACCAGCAG
AACACCCCCATCGGCGACGGCCCCGTGCTGCTGCCCGACAACCACTACCTGAGCACCCAGTCCGTGC
TGAGCAAAGACCCCAACGAGAAGCGCGATCACATGGTCCTGCTGGAGTTCGTGACCGCCGCCGGGA
TCACTCACGGCATGGACGAGCTGTACAAGTAAGCGGCCGCGACTCTAGATCATAATCAGCCATACCA
CATTTGTAGAGGTTTTACTTGCTTTAAAAAACCTCCCACACCTCCCCCTGAACCTGAAACATAAAATG
AA

### Supporting Tables

| FP | $\lambda_{max}$ (nm) | ε ($mM^{-1}cm^{-1}$) | $\lambda_{EM}$ (nm) | QY | Brightness ($mM^{-1}cm^{-1}$) | Relative photobleaching resistance[d] |
|---|---|---|---|---|---|---|
| EGFP[a] | 488 | 56 | 508 | 0.67 | 37.5 | 0.81 |
| sfGFP[b] | 485 | 49 | 511 | 0.72 | 35.3 | 1 |
| mNeonGreen[c] | 504 | 113 | 517 | 0.80 | 90.4 | 0.72 |
| mClover[c] | 505 | 105 | 516 | 0.84 | 88.2 | 0.27 |
| mVenus[c] | 515 | 127 | 528 | 0.67 | 85.1 | 0.13 |
| mCherry[c] | 586 | 85 | 610 | 0.30 | 25.5 | 1.41 |

**Table S1. Comparison of different fluorescent proteins.**
[a], from FPBase (https://www.fpbase.org) [1].
[b], from Reddington et al [2].
[c], from the Cranfill et al [3]
[d], based on t (1/2) values published by Cranfill et al [3]. Values relative to sfGFP, which is set to 1.

**Table S2.** H-bond frequency between chromophore and residue 148 over 10ns simulations

| Variant | H-bonds = 0 | H-bonds = 1 | H-bonds = 2 | % time H-bonding |
|---|---|---|---|---|
| sfGFP WT (H148) | 952 | 49 | 0 | 4.9 |
| sfGFP H148S | 512 | 409 | 80 | 48.9 |
| sfGFP H148T | 993 | 8 | 0 | 0.8 |
| sfGFP H148N [a] | 222 | 779 | 0 | 77.9 |
| sfGFP H148N [b] | 992 | 9 | 0 | 0.9 |
| sfGFP H148C | 977 | 24 | 0 | 2.4 |
| sfGFP H148A | 1001 | 0 | 0 | 0 |

a, sfGFP H148N where starting rotamermic form has the NH group of the carboxamide group is closest to the chromophore phenol O.
b, sfGFP H148N where starting rotamermic form has the O group of the carboxamide group is closest to the chromophore phenol O.

**Table S3.** H-bond frequency between chromophore and W1 water over 10ns simulations

| Variant | H-bonds = 0 | H-bonds = 1 | H-bonds = 2 | % time H-bonding | Time before distance > 0.4 nm[c] |
|---|---|---|---|---|---|
| sfGFP WT (H148) | 945 | 56 | 0 | 5.6 | 0.22 ns |
| sfGFP H148S | 805 | 196 | 0 | 19.6 | 2.24 ns |
| sfGFP H148T | 870 | 131 | 0 | 13.1 | 1.4 ns |
| sfGFP H148N [a] | 946 | 55 | 0 | 5.5 | 0.83 ns |
| sfGFP H148N [b] | 931 | 70 | 0 | 7 | 0.1 ns |
| sfGFP H148C | 973 | 28 | 0 | 2.8 | 0.53 ns |
| sfGFP H148A | 775 | 224 | 2 | 22.6 | 1.05 ns |

a, sfGFP H148N where starting rotamermic form has the NH group of the carboxamide group is closest to the chromophore phenol O.

b, sfGFP H148N where starting rotamermic form has the O group of the carboxamide group is closest to the chromophore phenol O.

c, refers to distance between the chromophore phenol O atom and the O atom of W1. Time is based 0.4 nm distance threshold broken for at least 0.2 ns.

**Table S4.** Spectral properties of sfGFP-H148X variants.

| FP | $\lambda_{max}$ (nm) | ε ($mM^{-1}cm^{-1}$) | $\lambda_{EM}$ (nm) | QY | Brightness ($mM^{-1}cm^{-1}$) |
|---|---|---|---|---|---|
| sfGFP[a] | 485 | 49.0 | 509 | 0.72 | 35.3 |
| $sfGFP^{H148C}$ | 398<br>498 | 34.4<br>11.1 | 513 | 0.32<br>0.50 | 11.0<br>5.6 |
| $sfGFP^{H148N}$ | 487 | 52.6 | 509 | 0.37 | 19.5 |

a, originally reported by Reddington et al. [2]

**Table S5.** Residency time of water molecules close to the chromophore.

| Water | sfGFP Sim 1 | sfGFP Sim 2 | sfGFP Sim 3 | sfGFP Average | YuzuFP Sim 1 | YuzuFP Sim 2 | YuzuFP Sim 3 | YuzuFP Average |
|---|---|---|---|---|---|---|---|---|
| | Residency time (ns)[a] | | | | | | | |
| W1 | 3.36 | 306.55 (0.22)[c] | 0.89 | **103.6 (1.49)** | 14.79 | 8.15 | 2.64 | **8.53** |
| W2 | 171.46 | 124.14 | 119.19 | **138.26** | 500[b] | 63.83 | 500[b] | **354.61** |
| W3 | 33.54 | 200.19 | 500[b] | **244.58** | 89.69 | 248.41 | 11.62 | **116.57** |
| W4 | 50.77 | 188 | 119.19 | **119.32** | 37.14 | 162.39 | 2.88 | **67.47** |
| W5 | 41.51 | 221.98 | 68.77 | **110.75** | 25.56 | 500[b] | 42.15 | **189.24** |
| W6 | 66.3 | 89.21 | 500[b] | **218.50** | 73.43 | 500[b] | 161.45 | **244.96** |

a, water residency time close to the chromophore a determined by a distance consistency (0.1 ns) above a distance of 1.5 nm from the β-methylene bridge atom.

b, retained residency (see definition above) over the course of the 500 ns simulation. The average is thus calculated with these values set to 500 ns.

c, W1 in sfGFP Sim 2 moves from it's original position to more internal position close to the chromophore. W1 becomes internalised within 0.22 ns (see Figure S11).

| Mutation | Primer sequence (5’ to 3’) |
|---|---|
| H148S | CTGGAATATAATTTCAACAGC**AGT**AATG<br>TTTATGACCCAGAATGTTGCCATC |
| H148C | CGATAAACAGAAAAATGGCATCAAAGCG<br>GCGGTAATATACACATT**GCA**GCTGTTG |
| H148A | GGAATATAATTTCAACAGC**GCT**AATG<br>AGTTTATGACCCAGAATGTTGCCATCT |
| H148N | TTTCAACAGC**AAC**AATGTGTATATTAC<br>TTATATTCCAGTTTATGACCC |
| H148S (Mammalian constructs) | CAACTTCAACAGC**AGC**AACGTCTATATCAC<br>TACTCCAGCTTGTGCCCCAGGATGT |

**Table S6.** Primer combinations and sequences for introducing H148X mutations. Bases in bold and underlined represent the mutation. Top primer is the “forward” and bottom the “reverse” primer in a whole plasmid PCR.

**Supporting Figures.**

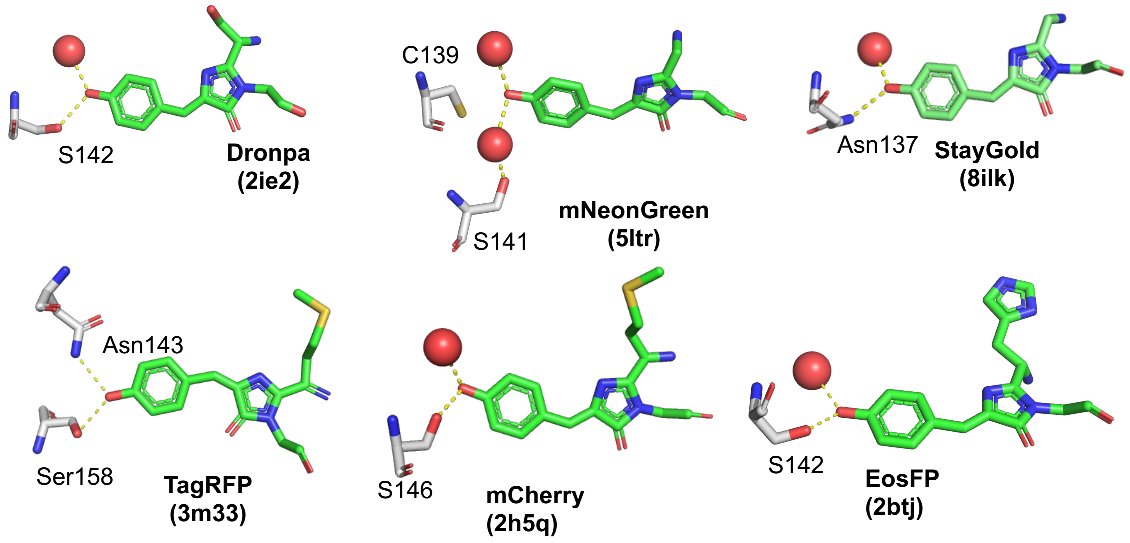

**Figure S1.** Additional examples of residues equivalent to H148 that interact with the chromophore phenol group: Dronpa [4] (PDB 2ie2), mNeonGreen [5](PDB 5ltr), StayGold[6] (PDB 8ilk), TagRFP [7,8] (PDB 3m33), mCherry [9](PDB 2h5q), EosFP [10] (PDB 1zux). In all cases the chomophore is coloured green, interacting residues equivalent to H148 in sfGFP grey and water molecules are red spheres.

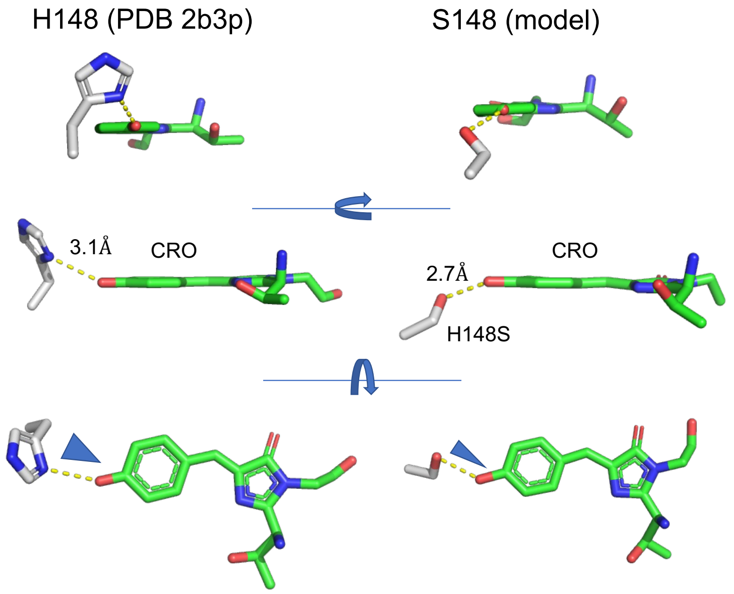

**Figure S2.** Relative conformations of residue 148 (grey sticks) and chromophore (green sticks). Left hand side is the position of H148 taken from the known structure of sfGFP (2b3p.pdb [11]). Modelled structure of the H148S is the clustered average of individual MD trajectory outputs after 10 ns of molecular dynamics.

H148 to:

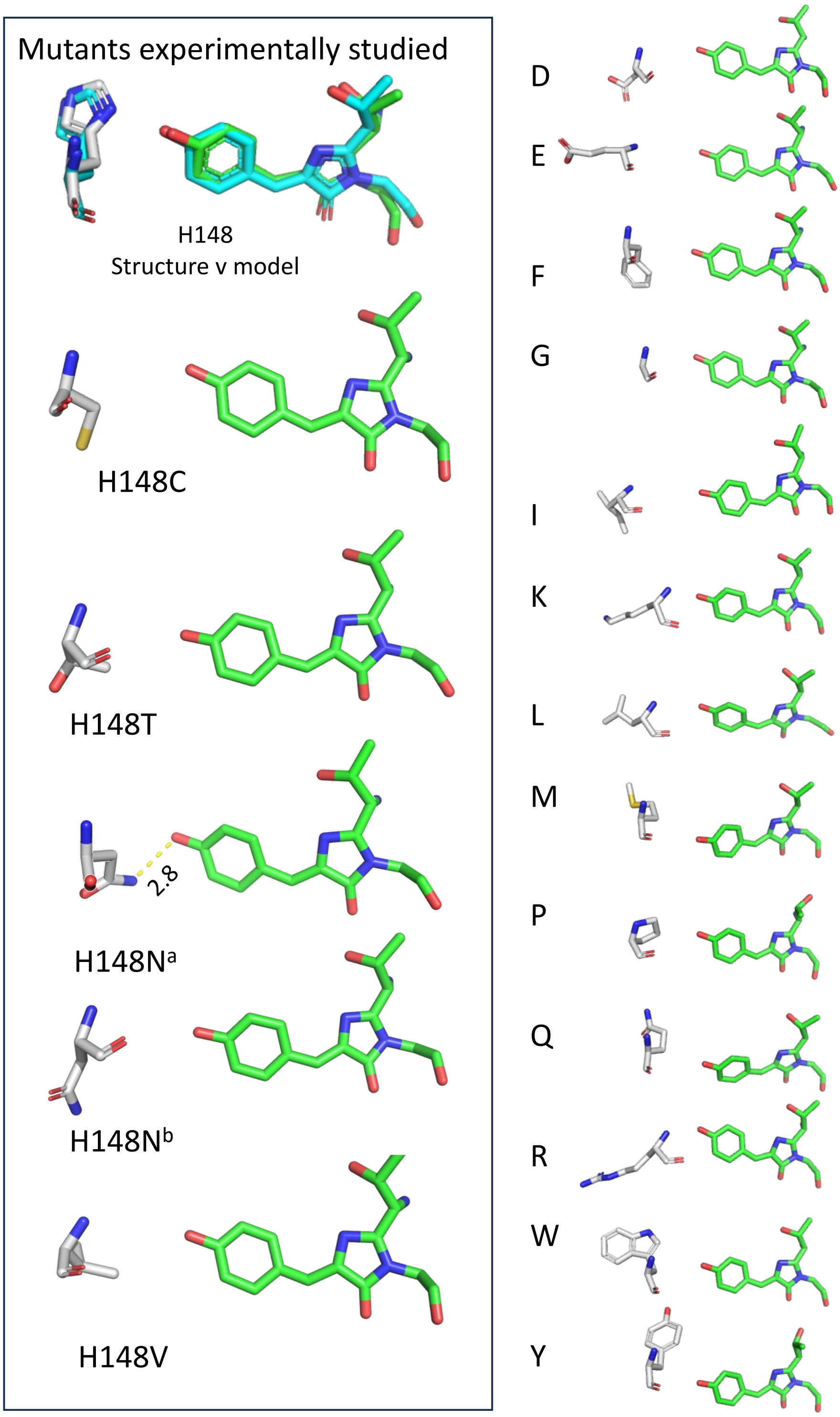

**Figure S3.** Modelling the H148X mutations. Modelled structures of the H148X mutations are the clustered average of individual MD trajectory outputs after 10 ns of molecular dynamics. Mutations on the left hand side have been experimentally analysed. For H148N, two model outcomes are shown: H148N[a] is from a starting point of the NH group of the carboxamide group closest to the chromophore phenol O; H148N[b] is from a starting of the O group of the carboxamide group closest to the chromophore phenol O.

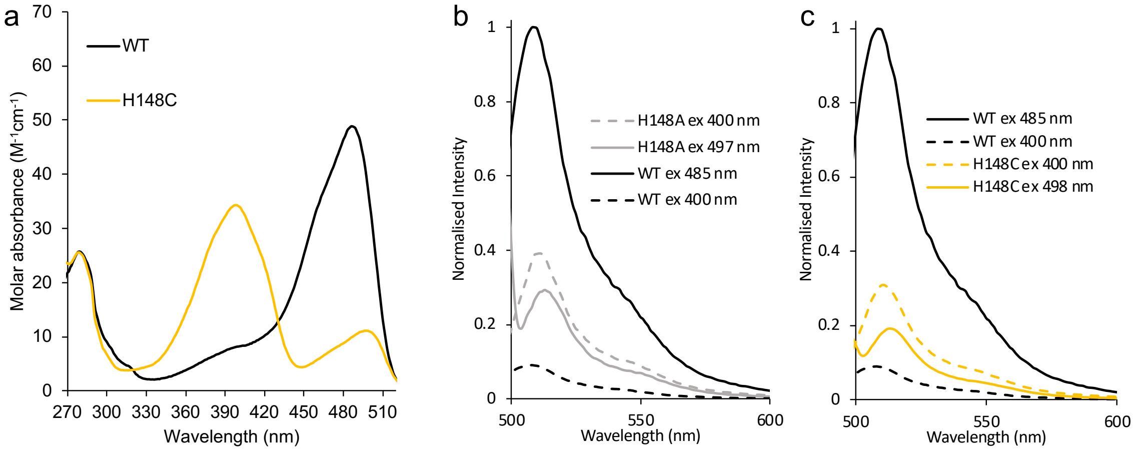

**Figure S4.** Absorbance and Fluorescence emission spectra for sfGFP variants. (a) Absorbance spectra for sfGFP WT (black) and H148C (orange). Fluorescence emission spectra for (b) H148A and (c) H148C. For clarity, the same WT sfGFP spectra are plot in each of the 4 panels and coloured black. Excitation at 485-498 nm is shown as solid lines and excitation at 400 nm as dashed lined. (b) H148A is coloured grey with excitation at 497 nm shown as solid lines and excitation at 400 nm as dashed lined. (c) H148C is coloured orange with excitation at 498 nm shown as solid lines and excitation at 400 nm as dashed lined.

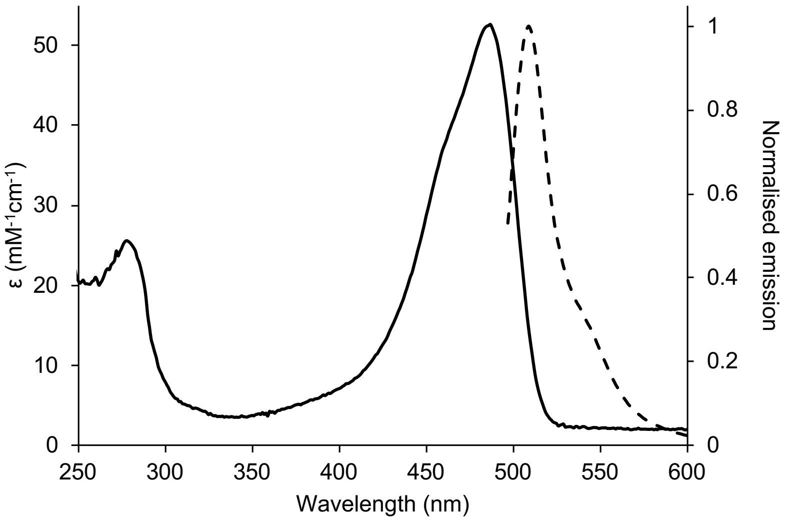

**Figure S5.** Absorbance and fluorescence spectra of sfGFP-H148N. The absorbance spectrum are shown as a solid line and normalised to the molar absorbance coefficient. Fluorescence emission spectra on excitation at 490 nm is shown as dashed lined and normalised to 1 based on the maximum emission intensity at 509 nm.

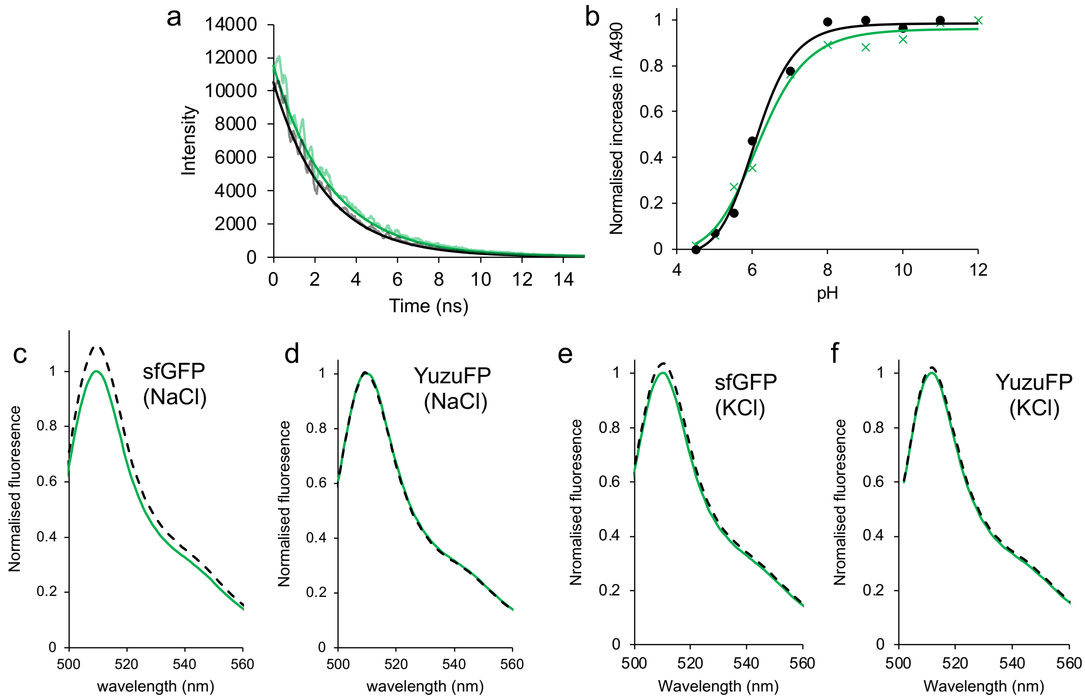

**Figure S6.** Analysis of YuzuFP. (a) The fluorescence intensity decay curve of YuzuFP (green) and sfGFP (black) on excitation at 473 nm. Data where fit to a single exponential decay in Graphpad Prism. (b) pKa fo YuzuFP (green) and sfGFP (black) as measured by the increase in absorbance at 490 nm. Data where fit to a least squares fit sigmodal model in Graphpad Prism. (c-f) The effect of 150 mM salt (either NaCl or KCl) on fluorescence emission, as indicated in the figure. Green solid lines represent protein in 50 mM Tris pH 8.0 and dashed black lines in the presence of 150 mM salt.

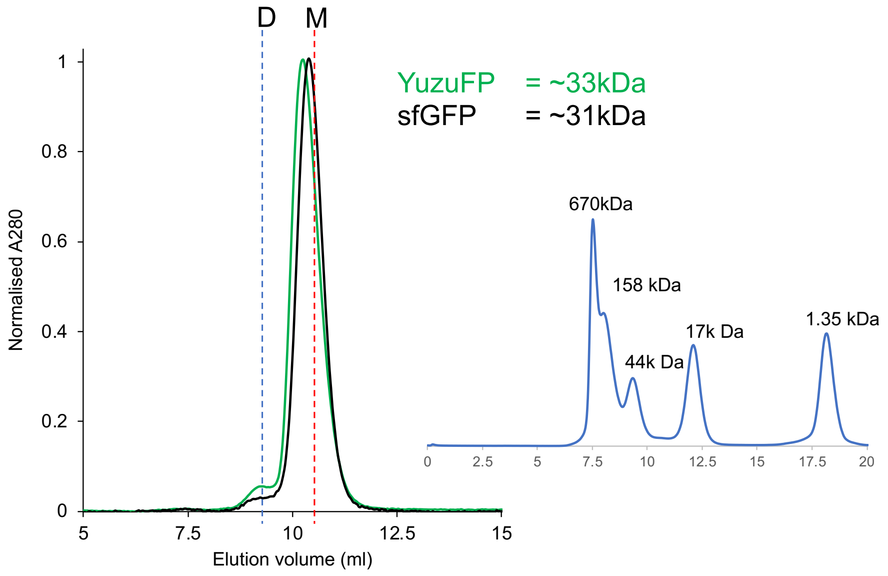

**Figure S7.** Analysis of quaternary structure YuzuFP by size exclusion chromatography (SEC). The black and green lines represent 50 μM sfGFP and 50 μM YuzuFP, respectively. SEC was performed using a calibrated Superdex 75 10/300GL column (inset). Estimated elution volumes for a theoretical monomer (M; 27 kDa) and dimer (D; 54 kDa) are shown by red and blue sashed lines, respectively. YuzuFP eluted very slightly earlier than sfGFP, giving an estimated molecular weight of ~33 kDa compared to 31 kDa for sfGFP. A small shoulder at ~9.3 ml suggests a small amount of dimer (~5%) may be present for both sfGFP and YuzuFP.

a.

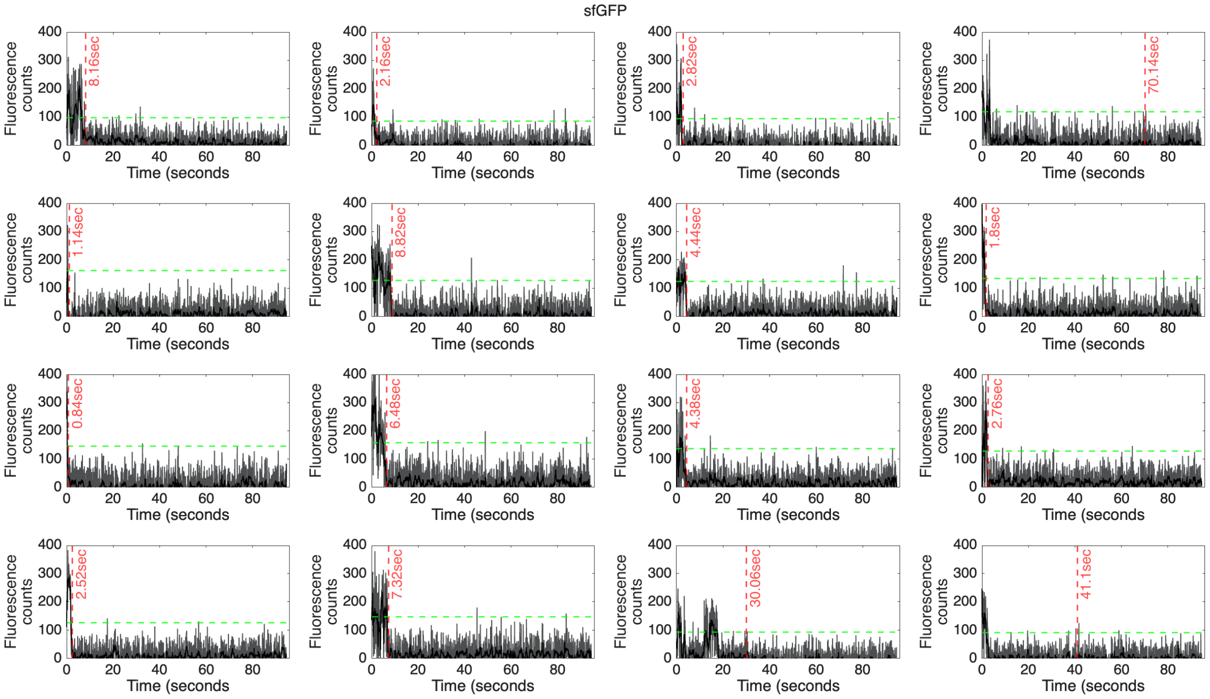

b.

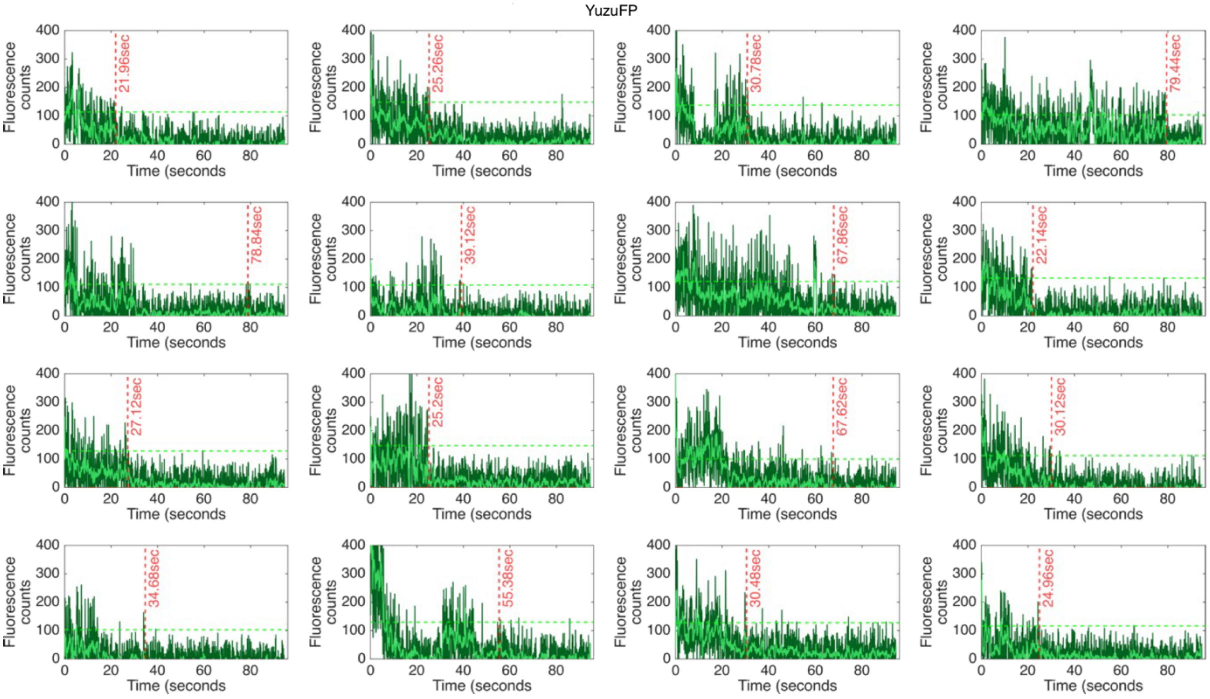

**Figure S8.** Exemplar single molecules fluorescence traces for (a) sfGFP and (b) YuzuFP . Single molecule data is extracted from a micrograph timeseries acquired using a total internal reflection fluorescence (TIRF) microscopy imaging system. Trace's are generated from the mean intensities of 4 by 4-pixel regions of interest corresponding to individual fluorophores. Raw data (grey and dark green) are plotted along side data passed through a forward-backward moving window average filter (black and light green). In order to clearly identify "on" and "off" fluorescence states thresholding of the raw data to 10 standard deviations beyond the mean background was used (green dashed line). Using this in combination with a temporal threshold allowed for identification of photobleaching lifetimes of all single molecules (red dashed lines and times in seconds).

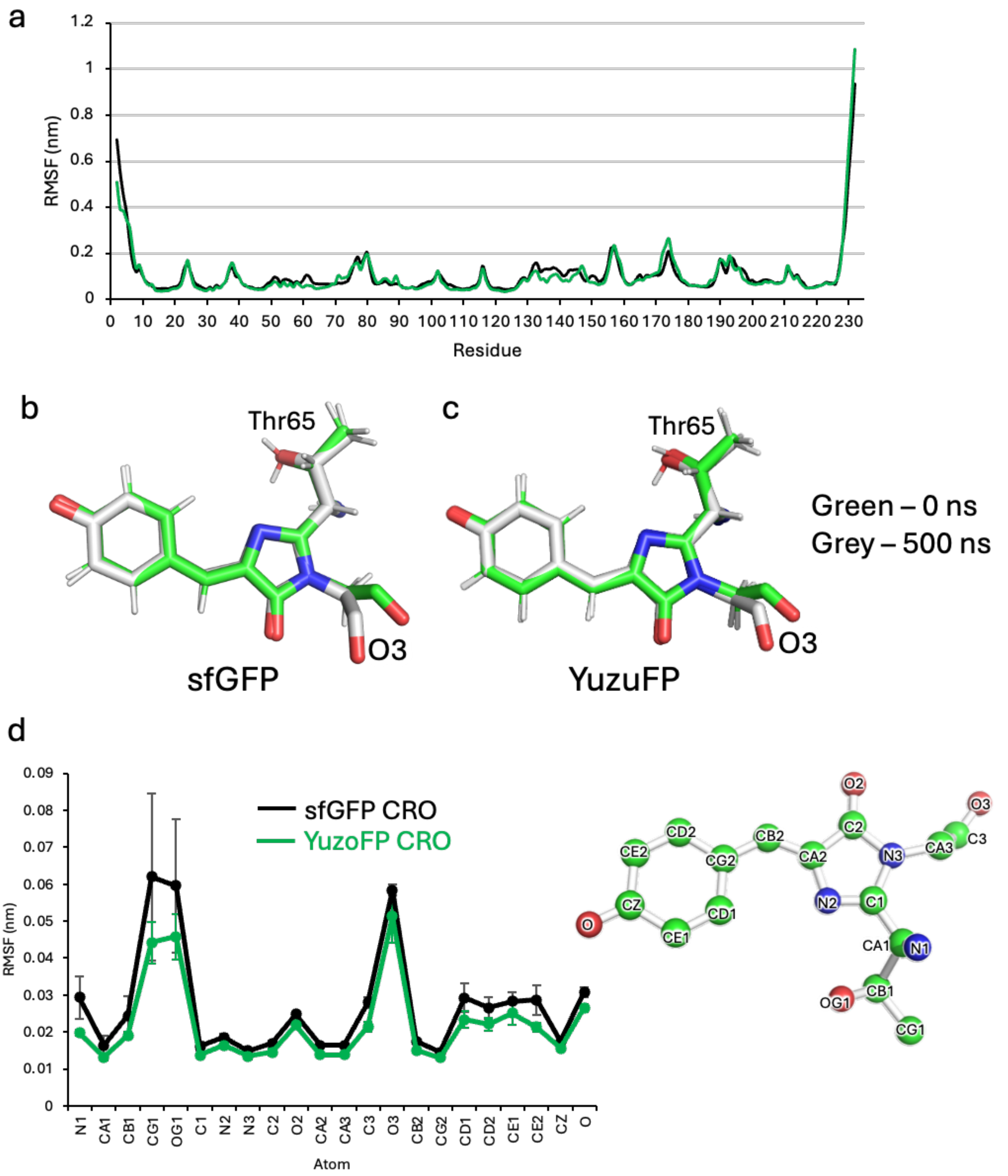

**Figure S9.** Change in backbone and chromophore structure over the course of MD simulation. (a) Per residue backbone (Cα) RMSF of sfGFP (black) and YuzuFP (green). RMSF values shown are an average of the 3 independent simulations. The chromophore structure at the start (green, 0 ns) and end (grey; 500 ns) of a single simulation for (b) sfGFP and (c)

YuzuFP. (d) The root mean square fluctuation (RMSF) of each of the chromophore heavy atoms (as annotated in the molecular structure to the right) for sfGFP (black) and YuzuFP (green). The error bars represent the standard deviation between the 3 independent 500 ns simula

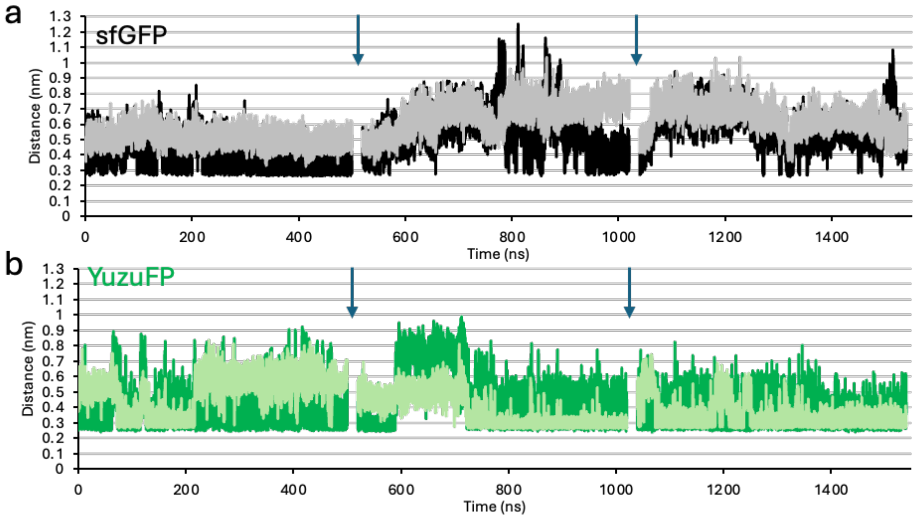

**Figure S10.** The pair-wise distance in (a) sfGFP and (b) YuzuFP, between chromophore phenol oxygen and residue 148 backbone and side chain H-bond donor heavy atoms. In (a), the backbone amide nitrogen is coloured grey and the side chain imidazole nitrogen of H148 is coloured black. In (b), the backbone amide nitrogen is coloured lime and the side chain hydroxyl oxygen of S148 is coloured green. The plots are a concatenation of the three 500 ns MD simulations, where the down arrows show the separation between each simulation.

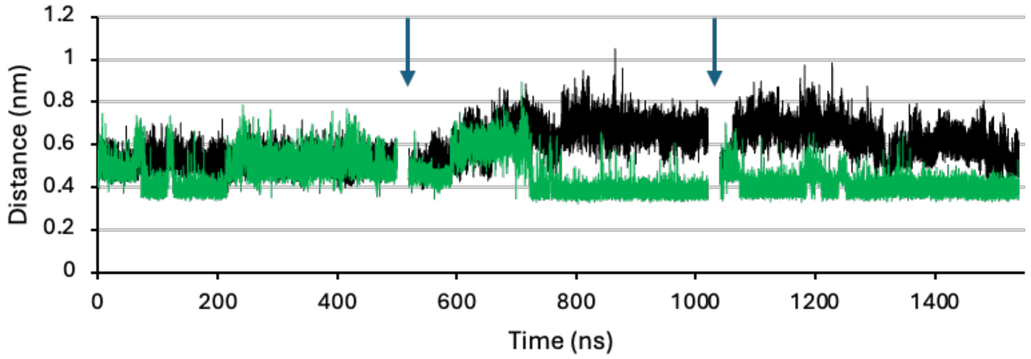

**Figure S11.** The distance between the chromophore phenol oxygen and the C$\alpha$ of residue 148 in sfGFP (black) and YuzuFP (green). Each plot is a concatenation of 3 separate 500 ns simulations. The start and end of each simulation is shown by the down arrows.

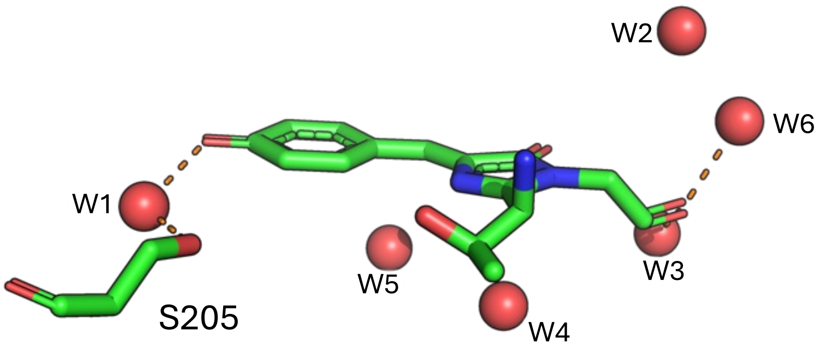

**Figure S12.** Water molecules within proximity of the chromophore. Red spheres are water molecules with the W1 water molecule labelled. Dashed lines are polar interactions between the chromophore and water molecules. The structure shown is PDB 2b3p.

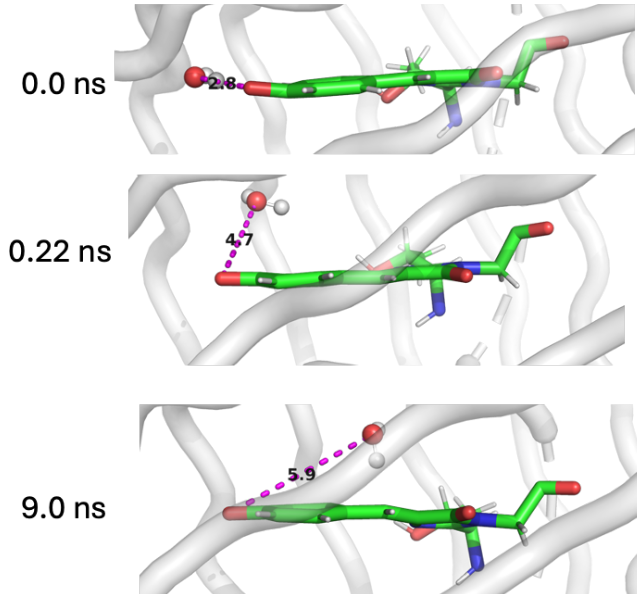

**Figure S13.** Position of water molecule W1 in sfGFP simulation 2. Within 0.22ns, W1 is internalised and no longer able to form a H-bond with the chromophore phenol oxygen. The chromophore is coloured green and water molecule W1 shown as balled and stick. The dashed magenta lines and associated numeric value represent the distance between the chromophore oxygen and the W1 water molecule (in Ångstroms).

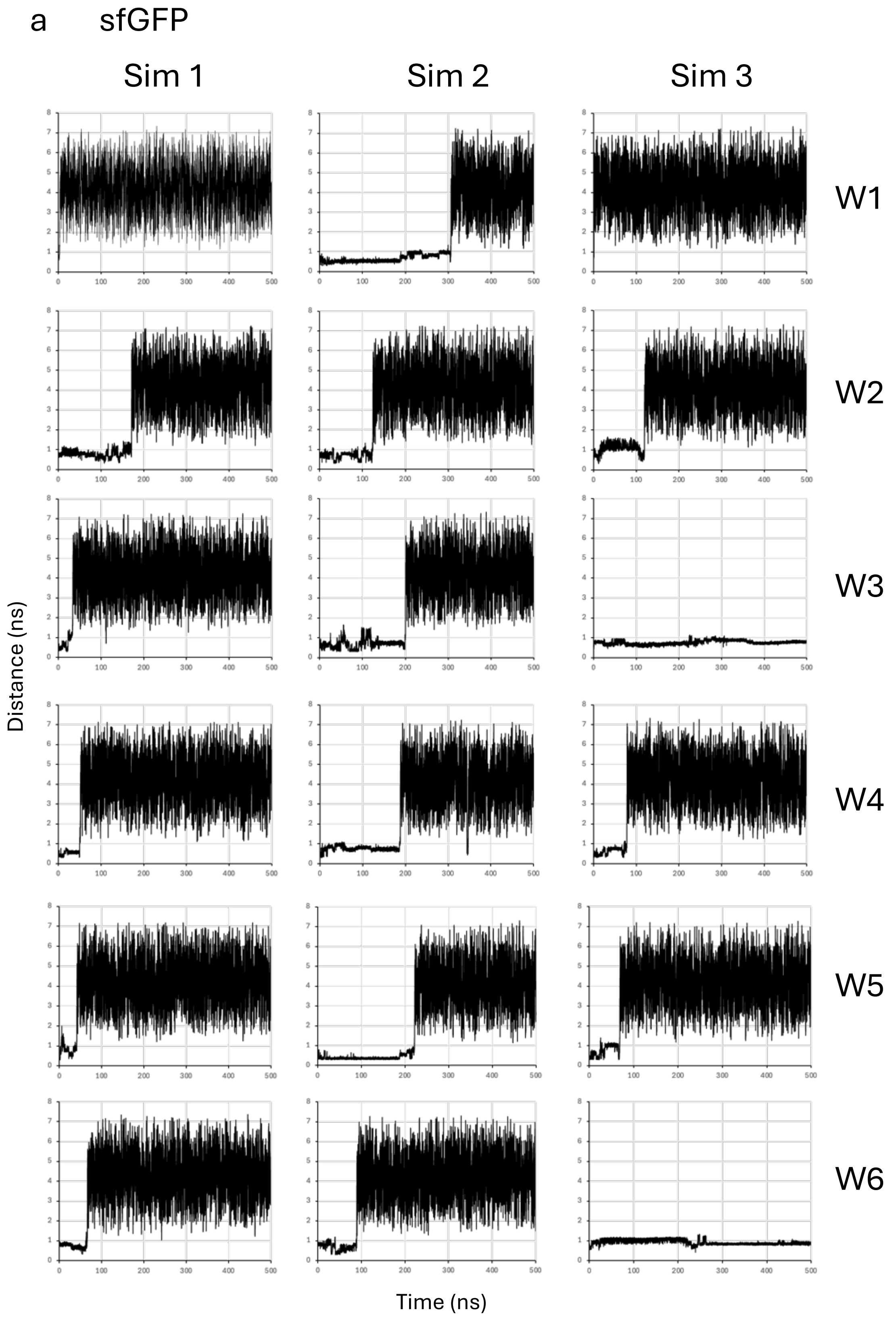

a
sfGFP
Sim 1
Sim 2
Sim 3
W1
W2
W3
W4
W5
W6
Distance (ns)
Time (ns)

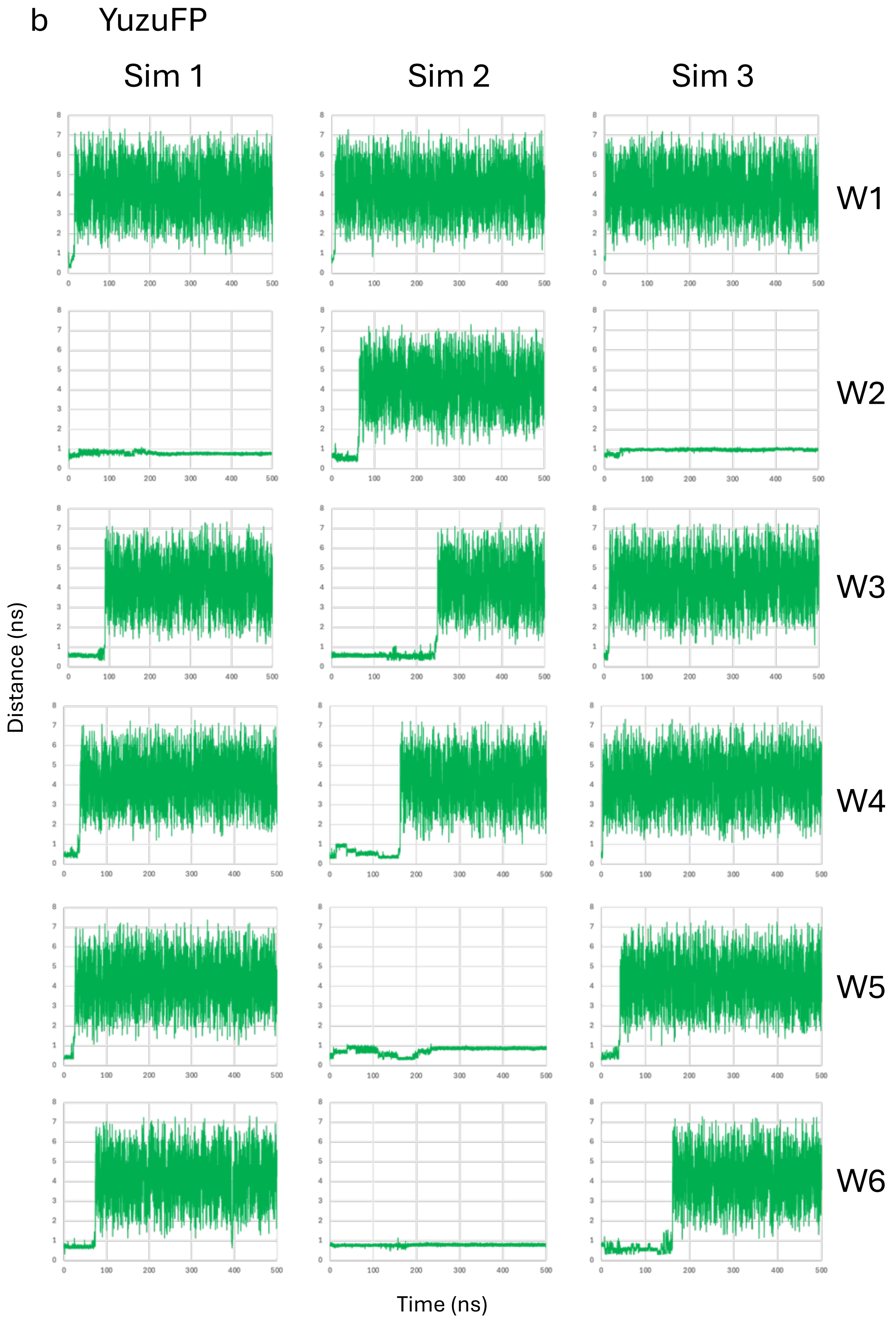

**Figure S14.** The pairwise distance between chromophore (atom 967 as part of the β-methylene bridge) and individual water oxygen atom (W1 to W6) from each 500 ns simulation (Sim 1 to 3) in (a) sfGFP and (b) YuzoFP. In sfGFP, W1 to W6 corresponding to oxygen atom IDs 3642, 3618, 3624, 3663, 3708, 3609, respectively. For YuzoFP W1 to W6 corresponding to oxygen atom IDs 3636, 3612, 3618, 3657, 3702, 3603.

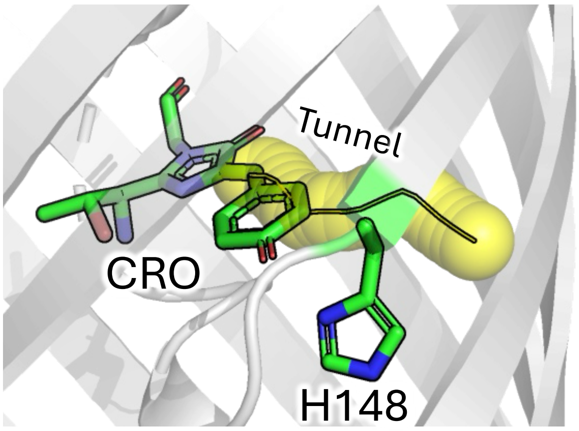

**Figure S15.** Tunnel (yellow; 1.1 Å probe radius) between CRO and solvent as calculated using CAVER [12,13] and the original PDB for sfGFP (2b3p)[11] .
